# Targeting the DNA damage repair protein RAD51 alters fibroblast metabolism and enhances apoptosis in pulmonary fibrosis

**DOI:** 10.64898/2026.04.01.715935

**Authors:** Rahul K. Maurya, Arun K. Sharma, Kyle J. Schaefbauer, Lina Ma, Jeffrey R. Koenitzer, Andrew H. Limper, Malay Choudhury

**Affiliations:** Division of Pulmonary and Critical Care Medicine, Department of Medicine, Washington University in St. Louis, MO 63110, USA; Department of Laboratory Medicine and Pathology, Mayo Clinic, Rochester, MN 55905, USA; Thoracic Disease Research Unit, Division of Pulmonary and Critical Care Medicine, Mayo Clinic, Rochester, MN 55905, USA

**Keywords:** Pulmonary fibrosis, DNA damage response, TGFβ signaling, RAD51, Oxidative phosphorylation, Glycolysis, Apoptosis, Precision cut lung slices, Bleomycin

## Abstract

**Background:** Idiopathic pulmonary fibrosis (IPF) is a progressive and fatal lung disease characterized by aberrantly activated, apoptosis-resistant profibrotic lung (myo)fibroblasts. Prior research has demonstrated that lung fibroblasts from patients with IPF exhibit resistance to DNA damage, suggesting that this behavior contributes to their persistent survival and continuous proliferation. We propose that elevated levels of the DNA damage repair protein RAD51 regulate myofibroblast activation and apoptosis and provide a potential therapeutic target to impede fibrosis progression.

**Methods:** Human lung fibroblasts were transfected with siRNA against RAD51 or treated with RAD51-specific inhibitor B02 and markers of fibrosis, DNA damage, apoptosis, metabolic reprogramming, and mitochondrial dynamics were assessed. The preclinical efficacy of B02 was evaluated in human precision cut lung slices (PCLS) and in a mouse model of pulmonary fibrosis.

**Findings:** RAD51 expression was significantly upregulated in the lungs and lung fibroblasts of IPF patients. Knockdown or inhibition of RAD51 in fibroblasts reduced profibrotic marker expression, suppressed mTORC1 signaling and mitochondrial function, and increased apoptosis susceptibility. Pharmacological inhibition of RAD51 shifted the profibrotic phenotype towards a fibrosis-resolving state in human and mouse PCLS, and in a bleomycin-induced mouse model of lung fibrosis.

**Interpretation:** The inhibition of RAD51 exerts therapeutic benefits in lung fibrosis by promoting apoptosis. Our findings identify that inhibiting RAD51 with B02 in fibroblasts impairs DNA repair and induces metabolic reprogramming, making it a potential therapeutic target.

**Research in context:** *Evidence before this study:* Pulmonary fibrosis (PF) is characterized by excessive fibroblast activation and subsequent deposition of extracellular matrix (ECM) proteins, which ultimately disrupt normal lung architecture. A significant contributing factor to the pathogenesis of pulmonary fibrosis is the presence of fibroblasts that are resistant to apoptosis, preventing normal wound healing. Recent studies highlight the DNA repair protein RAD51 as effective in protecting fibroblasts from death induced by chemotherapy and ionizing radiation. These finding suggested that RAD51 could have a role in fibroblast activation and apoptosis resistance in pulmonary fibrosis.

*Added value of this study:* We demonstrated that RAD51 is important for maintaining apoptosis-resistant fibrotic fibroblasts and their metabolic abnormalities. Our findings indicated that TGFβ-mediated upregulation of RAD51 reduces DNA damage, activates multiple pathways related to fibroblast activation and proliferation, and induces metabolic reprogramming, ultimately regulating apoptosis. Mechanistically, RAD51 inhibition enhanced p53 acetylation at lysine 120 and upregulated the expression proapoptotic proteins PUMA/BAK in mitochondria, promoting apoptosis. Pharmacological inhibition of RAD51 using the specific inhibitor B02 during the fibrotic phase of experimental lung disease effectively ameliorated pulmonary fibrosis.

*Implications of all the available evidence:* Our findings establish that RAD51 plays an important role in the survival of apoptosis-resistant fibrotic fibroblasts. We propose that reducing RAD51 expression leads to the metabolic reprogramming of activated fibroblasts, resulting in decreased mitochondrial respiration, reduced ATP levels, and diminished glycolysis or glutaminolysis. These observations suggest that targeting energy metabolism through RAD51 inhibition could be a viable strategy to enhance apoptosis, thereby creating a therapeutically targetable pathway in fibrotic cells. These findings highlight the potential of RAD51 as a therapeutic target for the treatment of IPF.

## Introduction

Idiopathic pulmonary fibrosis (IPF) is a chronic, progressive, and fatal interstitial lung disease of unknown etiology characterized by aberrant accumulation of extracellular matrix proteins, leading to distortion of the pulmonary architecture and impaired gas exchange.^1,2^ The worldwide incidence of IPF is estimated to range between 14 and 43 cases per 100,000 individuals with a median survival of 3-5 years from the time of diagnosis.^3,4^ Currently, three approved anti-fibrotic medications, pirfenidone, nintedanib and nerandomilast have been demonstrated to slow the decline in lung function in IPF patients, though they do not prevent disease progression or reverse existing fibrosis.^5–7^ These drugs all influence fibroblast function, suggesting that continued investigation of molecular mechanisms related to aberrant fibroblast activity is a valuable direction.

The accumulation of apoptosis-resistant myofibroblasts within fibrotic foci is a well-established phenomenon contributing to the pathogenesis of IPF.^8–10^ Compared to normal fibroblasts, those derived from the lungs of individuals with IPF exhibit increased survival rates when exposed to DNA-damaging chemotherapeutic agents such as cisplatin, potentially due to the upregulation of DNA repair proteins.^11^ RAD51, a key protein involved in the homologous recombination (HR) repair of DNA double-strand breaks, is activated by the Forkhead Box M1 (FoxM1) transcription factor. FoxM1 enhances the survival of lung fibroblasts from radiation-induced cell death by upregulating RAD51 and its associated repair pathways.^12^ Reduction of RAD51 levels promotes cellular senescence in alveolar epithelial cells, contributing to a cycle of aberrant healing.^13^ Increased activity or dependence on RAD51 in various cancers create therapeutic vulnerabilities when HR is inhibited. The small molecule inhibitor B02, which specifically targets RAD51, has demonstrated preclinical efficacy in both HR-deficient and HR-proficient tumors by reducing the formation of RAD51 foci, impairing HR, and sensitizing tumor cells to DNA-damaging agents and PARP inhibitors.^14–17^ Therefore, alterations in RAD51 activity, driven by various regulatory mechanisms, can significantly impact the pathogenesis of pulmonary fibrosis.

IPF fibroblasts maintain an apoptosis-resistant phenotype characterized by fewer DNA double-stranded breaks compared to non-IPF lung fibroblasts.^12^ Enhanced DNA repair mechanisms are important for maintaining the survival of these death-resistant fibrotic fibroblasts, thereby contributing to the progression of fibrosis.^12^ Here, we report that RAD51 inhibition or knockdown exhibits antifibrotic activity by modulating mTORC1 and p53 activity and enhancing the expression of anti-apopotic proteins PUMA/BAK, thereby promoting cell death. We found that RAD51 depletion inhibits glucose and glutamine transport, as well as mitochondrial oxidative phosphorylation in line with the emerging concept that targeting mitochondrial dysfunction and fibroblast metabolism to selectively eliminate metabolically defective and apoptosis-resistant fibrotic fibroblasts.^18,19^ These findings suggest that the metabolic changes induced by RAD51 could be therapeutically exploited as a promising strategy for the treatment of IPF.

## Materials and Methods

### Cell culture

Primary normal human lung fibroblasts (NHLF) and lung fibroblasts from patients with IPF were isolated from deidentified normal and IPF lung tissues. These cells were cultured in Dulbecco’s modified eagle medium glutaMax (DMEM; Gibco), supplemented with 10% fetal bovine serum (FBS; Hyclone Laboratories) and 1% penicillin-streptomycin (P/S; Life Technologies) at 37 °C in a humidified atmosphere containing 5% CO_2_. De-identified NHLF and IPF fibroblasts were obtained from Dr. Carol Feghali-Bostwick at the Medical University of South Carolina in Charleston, SC (University of Pittsburgh IRB #970946). Additionally, lung samples were obtained from consenting patients at the time of lung transplantation from IPF patients or non-transplantable lung tissues from deceased non-diseased donors at Barnes-Jewish Hospital at Washington University for the purpose of isolating fibroblasts, conducting precision-cut lung slice (PCLS) experiments, and performing immunohistochemistry or immunofluorescence staining (Washington University IRB #201103213). A summary of the pharmacological inhibitors utilized is provided in Table S1.

### Mice

Wild-type male and female C57BL/6 mice were purchased from Charles River Laboratories and used at 10 weeks of age. All studies involving mice were conducted in accordance with the Mayo Clinic-approved IACUC protocol #A00005741-20.

### Immunohistochemistry (IHC)

Formalin-fixed, paraffin-embedded lung tissue sections from both non-diseased and IPF lungs were used for IHC analysis. Antigen retrieval was performed on the paraffin-embedded lung tissue sections by heating the slides in citrate buffer (pH 6.0) using an antigen unmasking solution (Vector Laboratories) for 3 min, followed by a cooling period of 30 min. To block endogenous peroxidase activity and prevent non-specific antibody binding, the tissues were incubated with Bloxall (Vector Laboratories) and subsequently with 2.5% horse serum. The sections were then incubated overnight at 4 °C in a humidified chamber with primary antibodies against RAD51. This was followed by a 30-min incubation at room temperature with ImmPress polymer-bound horse anti-goat secondary antibodies (Vector Laboratories). The slides were stained with 3,3-diaminobenzidine (DAB, Vector Laboratories) for 1 min and counterstained with hematoxylin (Vector Laboratories) for 3 min.

### Immunofluorescence (IF) staining

NHLF cells (1x10^5^) were incubated on coverslips placed in 6-well plates containing 10% FBS/DMEM for 24 h. After this incubation period, the medium was replaced with 0.1% FBS/DMEM and treated with either vehicle (4 mM HCl/0.1% BSA) or TGFβ (5 ng/ml), with or without B02 (10 μM) for an additional 24 h. Cells cultured on coverslips, as well as lung sections from both normal and IPF lungs, along with saline and bleomycin-treated mouse lungs or human and mouse precision-cut lung slices (hPCLS or mPCLS), were fixed in 4% paraformaldehyde. They were subsequently permeabilized with 0.3% Triton X-100, blocked with 2% donkey serum, and incubated overnight at 4 °C with primary antibodies targeting ACTA2, CTHRC1, COL1, cleaved Caspase 3, p53K120, γH2AX and RAD51. Following three PBS washes, samples were incubated for 1 h at room temperature with fluorescently labeled secondary antibodies conjugated to Alexa Fluor 488 or Alexa Fluor 594 (Abcam), along with DAPI. Fluorescence images were acquired using a laser scanning confocal microscope (zeiss LSM 880 confocal laser scanning microscope) and analyzed for density using ImageJ software from NIH.

### Western blot analysis

Cells or lung tissues were lysed in ice-cold RIPA buffer (ThermoFisher Scientific), containing protease inhibitor mix (cOmplete, Roche) and 1 mM PMSF. The lysate was then centrifuged at 13,000xg for 15 min at 4 °C, and the supernatant was collected. Equal quantities of protein (5-20 μg) were thoroughly mixed with 5XSDS loading buffer, boiled for 5 min, and subsequently separated by 10-14% SDS-polyacrylamide gel electrophoresis. The resolved protein samples were transferred to an Immobilon-P PVDF Membrane (Millipore Sigma). The membranes were then incubated with primary antibodies (as specified in Table S1) overnight at 4 °C. Immunoreactivity was detected using enhanced chemiluminescence (ECL, ThermoFisher Scientific), and quantification was performed using ImageJ software from NIH.

### Real time quantitative PCR (qRT-PCR) analysis

Total RNA was isolated from cells using the RNeasy Plus Mini Kit (Qiagen). One microgram (1 μg) of RNA was reverse transcribed into cDNA using random primers (Life Technologies) and Maxima Reverse Transcriptase (Thermo Fisher Scientific). The resulting cDNA (equivalent to 1-5 ng of RNA) was subjected to real-time PCR amplification using the PowerTrack™ SYBR Green Master Mix (ThermoFisher Scientific) on a QuantStudio 3 Real-Time PCR System (Applied Biosystems). TATA box binding protein (TBP) was used as a normalization control. The relative expression levels of the target genes were calculated using the 2^-ΔΔCt^ method. The human and murine primer sets are provided in Table S2. All experiments were conducted with biological triplicates.

To isolate RNA from mouse lung tissue, the samples were lysed and homogenized using RLT Plus Buffer (RNeasy Plus Mini Kit; Qiagen). The lysate was then passed through a gDNA eliminator spin column to remove genomic DNA. Ethanol was added to the flow-through, and the samples were subsequently processed through a RNeasy Min elute spin column according to the manufacturer’s instructions.

### siRNA mediated gene knockdown

NHLF or IPF fibroblasts were transiently transfected with 40 nM small interfering RNA (siRNA) targeting RAD51, SMAD2, SMAD3, or with a non-targeting control siRNA (Santa Cruz Biotechnology), according to the manufacturer’s guidelines. The siRNAs from Santa Cruz Biotechnology were composed of three target-specific sequences. Briefly, 7.5x10^4^ cells were transfected using Lipofectamine 3000 (Invitrogen) and incubated in Opti-MEM (Invitrogen) for 6 h. Following this incubation period, the cells were transferred to complete medium (10% FBS/DMEM) for 18 h to facilitate recovery. Subsequently, the cells were treated with either vehicle or TGFβ in low serum medium (0.1% FBS/DMEM) for an additional 24 h. Following treatment, the cells were processed for western blot or qPCR analysis.

### Precision-cut lung slices (PCLS)

Adult non-diseased human or IPF lung tissue was inflated with 3% low melting point agarose (Invitrogen) to achieve uniform inflation. For mice treated with saline or bleomycin, lungs were inflated with 2% low melting point agarose. Subsequently, the lung tissue was placed in ice-cold Krebs bicarbonate buffer to facilitate agarose gel solidification and to ensure optimal tissue viability. The lung tissues were then embedded in a mold containing 6-8% low melting point agarose and stored at 4 °C for 5 min to allow polymerization. To create slices of 250 µm thickness, a vibrating microtome (Precisionary Instruments) was used. The resulting slices were collected and maintained in Hibernate-A medium (Gibco) at 4 °C for 15 min. Slices were then incubated in adult neurobasal media (Gibco) supplemented with 2% B27 and 1% penicillin-streptomycin (P/S) for 24 h. Following this incubation period, healthy slices were treated with 10 ng/ml of TGFβ and TNFα and incubated for an additional 72 h before proceeding to IF or western blotting. To assess the viability of the cultured slices, a WST-8 cell proliferation assay (Abcam) was performed. After the 72 h incubation, 100 µl/ml of WST-8 reagent was added to each well and incubated for 2 h. Absorbance was measured at 440 nm using a microplate reader. Wells containing only medium and WST-8 without tissue slices served as background controls.

### Metabolic function assay

NHLFs (2x10^5^ cells per well) were cultured in 6-well plates using DMEM complete medium (10% FBS, 1% Penicillin-Streptomycin) for 24 h. Following this, the cells were serum-starved (0.1% FBS, 1% Penicillin-Streptomycin in DMEM) for an additional 24 h. Subsequently, the cells were pretreated with either DMSO or B02 (10 µM) at 37 °C for 1 h, followed by treatment with either vehicle or TGFβ for 48 h. After treatment, the cells were seeded into Seahorse XF24 cell culture microplates (Agilent Technologies). The oxygen consumption rate (OCR) and extracellular acidification rate (ECAR) were measured using a Seahorse XFp extracellular flux analyzer (Agilent Technologies), employing the XFp cell mito stress test kit and the XF glycolysis stress test kit (Agilent Technologies).

### Measurement of glutamine

The intracellular level of glutamine was determined using the glutamine assay kit (BioVision) according to the manufacturer’s instructions. Briefly, NHLF cells were seeded in 6-well plates at a density of 2x10^5^ cells per well in a medium consisting of 10% FBS in DMEM and incubated for 24 h. Following this, the medium was replaced with 0.1% FBS in DMEM, and the cells were incubated for an additional 24 h. Subsequently, the cells were pretreated with either DMSO (0.1%) or B02 (10 μM) for 1 h, followed by stimulation with either vehicle or TGFβ (5 ng/ml) for 24 h and the glutamine levels were measured.

### Quantification of lactate level

Intracellular lactate levels were measured using the PicoProbe™ lactate fluorometric assay kit (BioVision). Briefly, NHLF cells were treated with TGFβ (5 ng/ml) and/or B02 (10 μM) for 48 h before being harvested. Subsequently, cell lysates (50 μl) were combined with 50 μl of a reaction mixture containing enzyme mix, PicoProbe reagent, and lactate assay buffer, and then incubated for 30 min at room temperature in the dark. Fluorescence was measured at Ex/Em = 535/587 nm using a microplate reader. Lactate concentration was determined from a standard curve and normalized to total protein content.

### ATP Assay

Intracellular ATP concentrations were quantified utilizing the ATP colorimetric/fluorometric assay kit (BioVision). In brief, cell lysates were prepared in ATP assay buffer, and 50 µl of each lysate was mixed with a reaction mixture containing an ATP converter, ATP probe, and developer. The mixture was incubated at room temperature for 30 min. Absorbance was measured at 570 nm, and ATP concentrations were calculated by referencing a standard curve and normalizing to protein levels.

### NADP^+^/NADPH measurement

The NADP+/NADPH ratio was determined using the NADP^+^/NADPH quantification kit (BioVision) following the manufacturer’s instructions. In brief, 2x10^5^ cells were pretreated with either DMSO (0.1%) or B02 (10 μM) for 1 h and subsequently stimulated with either vehicle or TGFβ (5 ng/ml) for 24 h. The cells were then collected in NADP/NADPH Extraction Buffer and underwent two freeze/thaw cycles, −80 °C to room temperature. The concentrations of NADP^+^ and NADPH were determined by comparing to standard curves and normalized to total protein content.

### The mitochondrial permeability transition pore (mPTP) opening assay

The mPTP opening assay was performed using the Image-IT™ live mitochondrial transition pore assay kit (ThermoFisher Scientific) following the manufacturer’s protocol. NHLFs were seeded at a density of 1x10^5^ cells per well in a 6-well plate, either in the presence or absence of TGFβ (5 ng/ml) with or without B02 for 24 h. After 24 h, the cells were incubated with 1 μM calcein-AM and 150 nM MitoTracker™ Red, with or without 1 mM CoCl_2_, in a modified Hanks’ Balanced Salt Solution (1x HBSS) for 15 min at 37 °C. Following incubation, cells were washed three times with 1x HBSS to remove excess dye. Cell imaging was performed using a Zeiss LSM 880 confocal laser scanning microscope, and fluorescence intensity was quantified using ImageJ software (NIH).

### Caspase 3 activity assay

The caspase 3 activity assay was performed using the EnzChek caspase-3 fluorescent assay kit (ThermoFisher Scientific) following the manufacturer’s instructions. NHLFs were seeded at a density of 6x10^3^ cells per well in 96-well optical-bottom black microplates (ThermoFisher Scientific) and incubated for 24 h. Subsequently, the cells were serum-starved for 24 h and treated with or without B02 in the presence or absence of TGFβ. After the treatment period, the cells were washed with 100 μl of ice-cold PBS and lysed on ice for 30 min. Subsequently, 50 μl of 2x reaction buffer containing 10 mM dithiothreitol and the caspase-3 substrate Z-DEVD-rhodamine 110 (5 μM) was added to each well. The plate was then incubated at 37 °C for 2 h. Caspase 3 activity was measured using a fluorescence microplate reader, and the results were expressed as relative fluorescent units (RFU).

### Bleomycin model of pulmonary fibrosis

Ten-week-old male and female C57BL/6 mice, maintained on breeder chow, were administered 50 μl of bleomycin (BLM; 2 U/kg body weight) or 0.9% normal saline via intratracheal instillation on day 0, under ketamine/xylazine anesthesia. On day 11, BLM- and saline-challenged animals were randomly assigned to receive intraperitoneal injections of either vehicle or B02 (50 mg/kg) every two days, for a total of six injections, and were euthanized on day 23. To evaluate respiratory function, the fur around the neck was shaved, and dissolved oxygen or oxygen saturation (SpO_2_) was measured using the MouseOx monitoring system every third day. On day 23, the animals were anesthetized, and flexiVent (Scireq) analysis was performed. Following euthanasia with pentobarbital (100 mg/kg), lungs were excised, weighed, and processed for hydroxyproline assay, histological analysis, and quantification of markers of fibrosis.

### Hydroxyproline assay

The total collagen content in the lungs was determined using the hydroxyproline assay kit (Millipore Sigma). Briefly, 20 mg of mouse lung tissue was homogenized in H_2_O at a concentration of 100 mg/ml. A 100 µl aliquot of the homogenate was then mixed with 100 µl of 12 M hydrochloric acid and hydrolyzed overnight at 120 °C. Following hydrolysis, a 10 µl aliquot of the resultant sample was subjected to colorimetric analysis for hydroxyproline, according to the manufacturer’s instructions.

### Histological Scoring

A blinded histologic scoring assay was performed to evaluate the progression and resolution of fibrosis in H&E-stained lung tissue from mice. This assessment involved a semiquantitative analysis that considered both the intensity and extent of fibrosis in each mouse lung, with the results being summed. The fibrosis scoring criteria were as follows: 0 (no fibrosis), 1 (focal or minimal fibrosis), 2 (multifocal or moderate fibrosis), and 3 (confluent or severe fibrosis).

### Statistics

Statistical significance were determined using GraphPad Prism 10.6 software. In vitro and ex vivo data were obtained from a minimum of 3 biological replicates. In vivo data consisted of 6-7 mice per group. Results are presented as the mean ± SEM. Comparisons between two groups were evaluated using a two-tailed unpaired Student’s t-test. For comparisons involving more than two groups, one-way or two-way ANOVA was performed, followed by Tukey’s post hoc analysis. Statistical significance is indicated by the number of asterisks: *P < 0.05, **P < 0.01, ***P < 0.001, ****P < 0.0001. A p-value of less than 0.05 considered significant.

## Results

### RAD51 is upregulated in pulmonary fibrosis and by TGFβ

Targeting DNA damage response (DDR) proteins could potentially increase the apoptotic susceptibility of fibroblasts, selectively removing them from fibrotic areas and alleviating fibrosis in patients with IPF. To investigate this hypothesis, we first analyzed DDR gene expression in the lungs of IPF patients. A microarray gene expression dataset from the Lung Genomics Research Consortium (LGRC), comprising 122 IPF and 91 control patients, revealed that DDR genes such as ATM, ATR, RAD51, BRCA1 and BRCA2, exhibited significantly upregulated mRNA expression in IPF patient lungs compared to control patients (**Fig. 1a, Fig. S1a**). To identify potential therapeutic targets, DDR proteins were screened with known inhibitory agents and examined their impact on TGFβ-mediated activation in Normal human lung fibroblast (NHLF). Inhibition of RAD51 most significantly reduced TGFβ-mediated induction of profibrotic molecules (**Fig. S1b**). Levels of RAD51 expression was also found to correlate with the severity of disease by pulmonary function, as measured by the percentage of predicted diffusing capacity for carbon monoxide (% DLCO) (**Fig. 1b**) and the percentage of predicted forced vital capacity (FVC; a measure of lung compliance) (**Fig. 1c**). Consistent with these observations, RAD51 protein expression was markedly elevated in lung tissues from patients with end-stage IPF undergoing lung transplantation, and colocalized with COL1^+^ (Collagen I) and ACTA2^+^ (Alpha smooth muscle actin) myofibroblasts (**Fig. 1d, e, and f, Fig. S1c**). Furthermore, fibroblasts isolated from the lungs of IPF patients exhibited significantly higher basal levels of RAD51 compared to NHLF (**Fig. 1g, h**). qPCR and western blot analyses further validated and extended these findings, demonstrating that TGFβ induces a time-dependent upregulation of RAD51 mRNA and protein expression in NHLF (**Fig. 1i and j, Fig. S1d**). This activation was specific to TGFβ stimulation, as treatment with the TβRI kinase inhibitor SB431542 completely abrogated the TGFβ-induced response in NHLF (**Fig. 1k, Fig. S1e**).

**Fig. 1:**
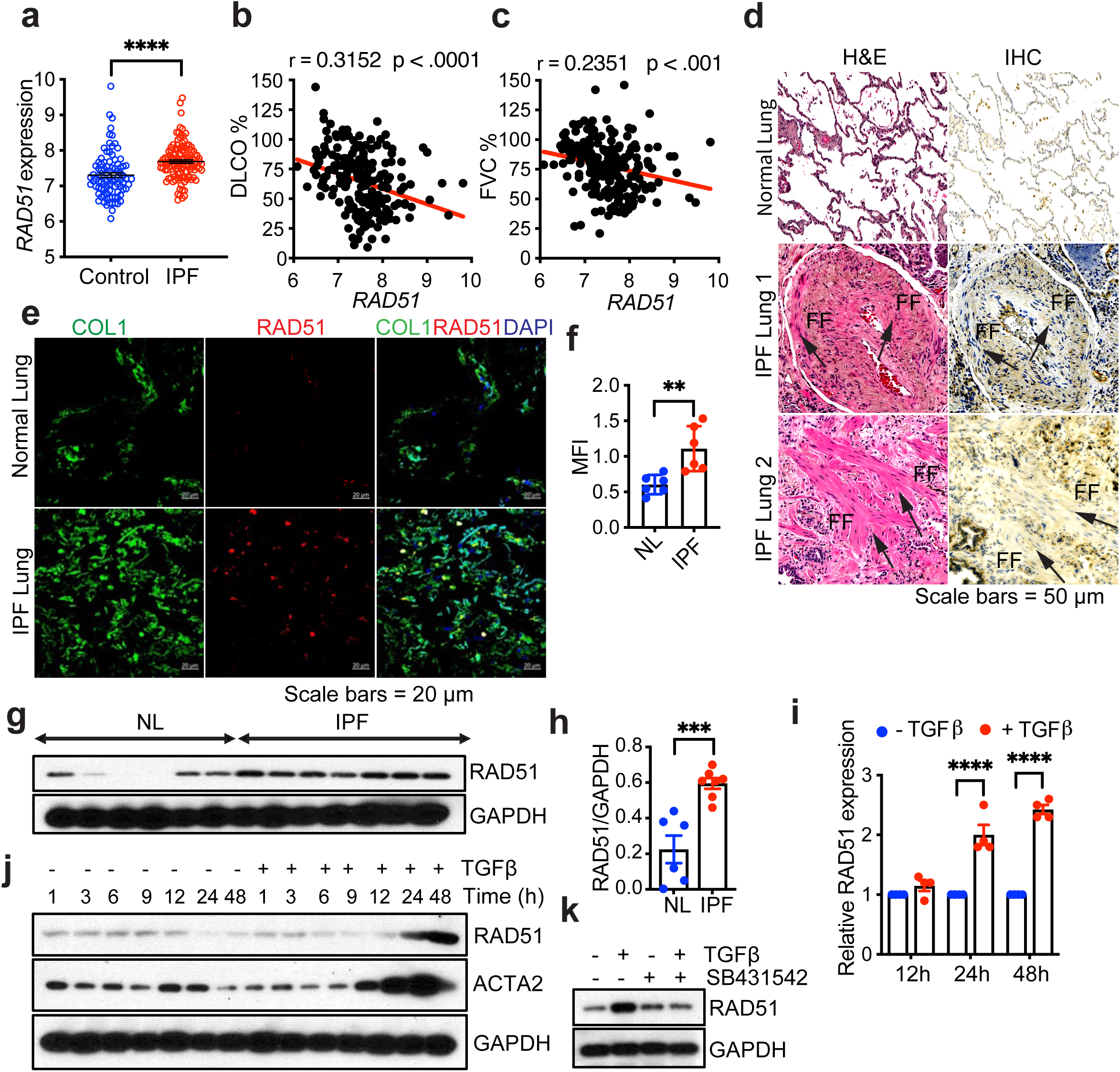
RAD51 is upregulated in pulmonary fibrosis and by TGFβ. **(a)** *RAD51* gene expression from 122 IPF and 91 controls patients of Lung Genomics Research Consortium (LGRC) microarray dataset (accession number GSE47460). **(b, c)** Correlation between *RAD51* expression and diffusing capacity of the lung for carbon monoxide (DLCO) or forced vital capacity (FVC) using the Pearson’s correlation coefficient (r). **(d)** Lung tissues from non-diseased normal (n=3) and IPF patients (n=3) were analyzed by H&E and IHC staining with anti-RAD51 antibody. Fibrotic foci (FF) are indicated in the IPF lung. Scale bars= 50 μm. (**e, f**) Localization of RAD51 was detected through immunofluorescence staining in lung tissue sections from control and IPF patients with antibodies for collagen 1 (green), RAD51 (red), and DAPI for nuclei (blue). Co-localization is observed in the merged images (far right side) for both non-IPF and IPF lung tissues. Scale bars= 20 μm. (**f**) Quantification of Mean Fluorescence Intensity (MFI) of RAD51 from (e). At least five random fields were analyzed through ImageJ program. (**g**) Primary lung fibroblasts from patients with IPF and healthy, non-diseased controls were cultured for 3-4 passages, followed by western blot analysis to assess RAD51 levels. (**h**) The ratios of RAD51 to GAPDH. (**i**) Normal Human Lung Fibroblasts (NHLF) cells were treated with 5 ng/ml TGFβ or a vehicle (4 mM HCL + 10 mg/ml BSA) for durations of 12 h, 24 h, and 48 h, followed by qPCR analysis of RAD51. (**j**) The western blot analysis of RAD51 was determined in quiescent NHLF cells, in the absence (-) and presence (+) of 5 ng/ml TGFβ, at the specified time points. (**k**) NHLF cells were subjected to treatment with DMSO (0.1%) or the TβRI inhibitor SB431542 (10 μM), either in the presence or absence of TGFβ (5 ng/ml, 48 h induction), and protein expression of RAD51 was determined by western blotting. Data generated for qRT-PCR represent mean ± SEM of n = 3 independent experiments. IHC, IF and western blots are representative of 3 independent experiments. Differences between groups were evaluated by two-way (i) ANOVA test with Tukey post-hoc analysis or unpaired two-tailed student’s t-test (a, b, c, f, h) using GraphPad Prism 10.6 software. ***P < 0.001, ****P < 0.0001.

### Profibrotic TGFβ signaling requires RAD51 and its inhibition decreases DNA repair in fibroblasts

We next investigated the role of RAD51 in TGFβ-mediated profibrotic signaling. B02, a selective inhibitor of the RAD51 protein, disrupts RAD51 DNA damage foci formation that accumulate at DNA double strand break sites, which are essential for homologous recombination and DNA repair.^20^ We performed a WST-1 cell proliferation assay to evaluate cellular toxicity, which demonstrated that B02 did not significantly affect cellular proliferation or exhibit toxicity at the 10 μM concentration used to impact profibrotic TGFβ signaling. (**Fig. S2a**). The expression of RAD51 was inhibited using siRNA or B02, and the impact on TGFβ-induced profibrotic marker expression was determined by western blotting and qPCR in NHLF (**Fig. 2a and b, Fig. S2b and c**) and IPF fibroblasts (**Fig. 2c and d**). Upon RAD51 inhibition or knockdown, the expression of profibrotic targets including COL1 (Collagen I), FN (Fibronectin), CTGF (Connective tissue growth factor), and ACTA2 (Alpha smooth muscle actin) were diminished. (**Fig. 2a and b, Fig. S2b and c, Fig. S3a-d).** Immunostaining revealed that TGFβ-treated NHLF cells exhibited pronounced ACTA2 and F-actin-positive stress fibers, which were reduced following B02 treatment (**Fig. 2e and f**). The functional role of RAD51 was further evaluated through a fibroblast migration assay induced by TGFβ. Cell migration, measured after wounding a confluent monolayer, was significantly reduced in the presence of B02 (**Fig 2g and h**). Collectively, these findings indicate that myofibroblast differentiation and function in part depend on RAD51 availability in TGFβ-mediated fibroblast activation. As an indirect marker of DNA double-stranded breaks (DSB), we next assessed the levels of phosphorylated H2AX (γH2AX), which forms descrete foci around DNA DSB sites to promote the accumulation of DNA repair proteins.^21,22^ Enhanced γH2AX staining was observed in both IPF fibroblast and NHLF cells treated with B02 (**Fig 2i-l**). Western blot analyses confirmed that TGFβ decreased γ expression, whereas sustained γH2AX expression was observed in RAD51-inhibited lung fibroblasts (**Fig. 2m, Fig. S3e**). These results suggested that inhibiting RAD51 activity increases DSBs, as demonstrated by the accumulation of γH2AX, thereby reducing the activation of fibroblasts.

**Fig. 2:**
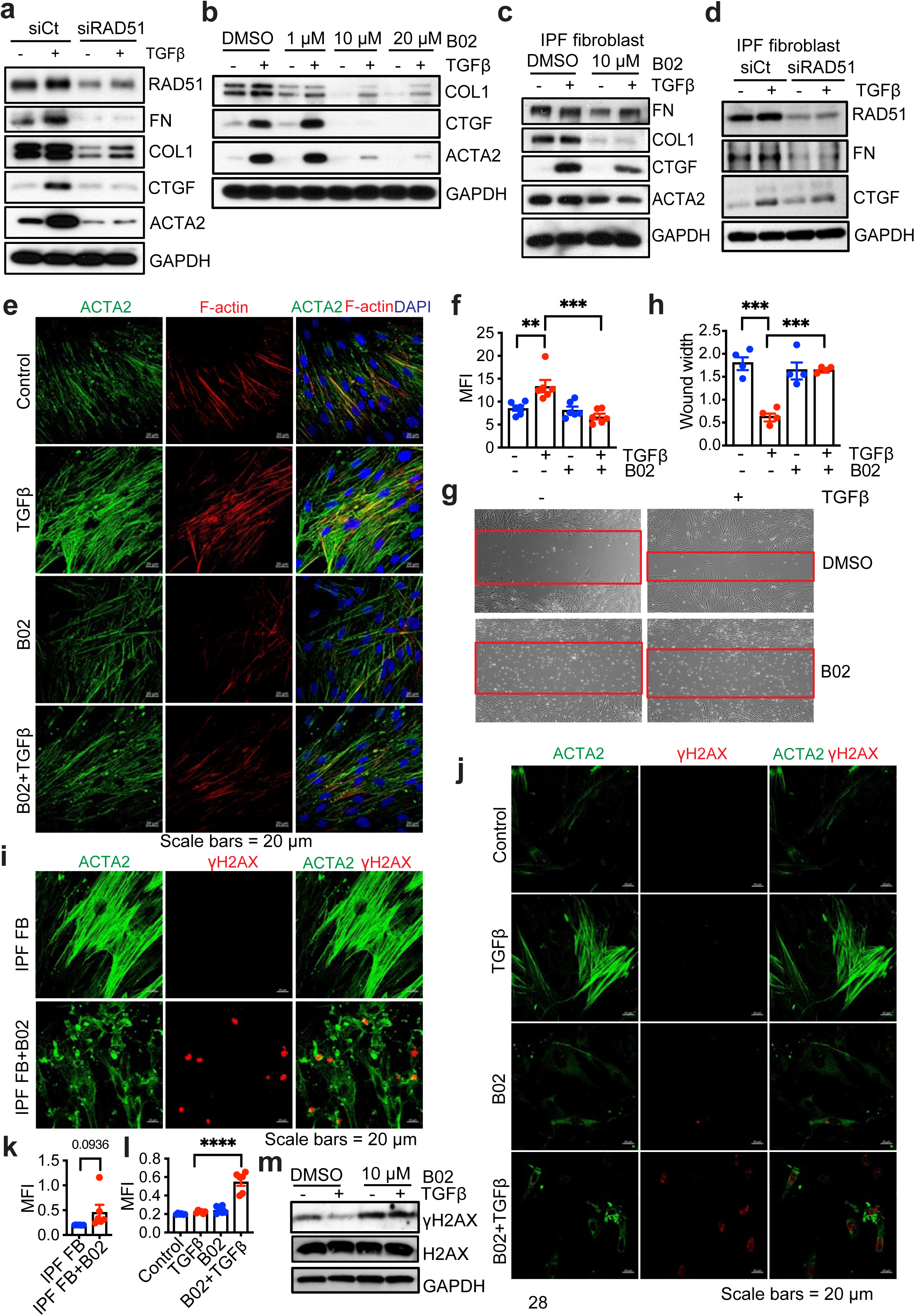
Profibrotic TGFβ signaling requires RAD51 and its inhibition decreases DNA repair in fibroblasts. (**a**) NHLF cells were transfected with either a non-targeting control (siCt) or siRNA targeting RAD51 (siRAD51). A vehicle (-) or TGFβ (+) was subsequently added to a final concentration of 5 ng/ml. After 48 h incubation period, western blotting analyses of the specified profibrotic molecules were performed. (**b**) Quiescent NHLF cells were pretreated for 1 h with 0.1% DMSO or 1, 10 and 20 μM of B02 (RAD51 specific inhibitor) prior to addition of vehicle or TGFβ (5 ng/ml). Following 48 h incubation, cell lysates were prepared and western blotted for indicated proteins. (**c, d**) IPF fibroblasts were either treated with vehicle (0.1% DMSO) or 10 μM B02 (c) or transfected with non-targeting control (siCt) or siRNA against RAD51 (d) and western blotted as mentioned in Fig. 2a and b. (**e**) Immunofluorescence staining of ACTA2 and F-actin in NHLF cells with/without 1h B02 (10 μM) pretreatment followed by addition of vehicle (-) or TGFβ (5 ng/ml) for 24 h. (**f**) Quantification of ACTA2 by mean fluorescence intensity (MFI) using ImageJ. (**g**) Scratch assays were performed on NHLF cells. The red bands signify the leading edge after 24 h in the presence (+) or absence (-) of TGFβ (5 ng/ml), either alone or in combination with B02 (10 μM), and are representative of 3 separate experiments. (**h**) Quantification of wound closure. Data reflect mean ± SEM of n = 3. (**i**) Immunofluorescence staining of γH2AX and ACTA2 in IPF fibroblasts with/without 24 h B02 (10 μM) treatment. (**j**) Immunofluorescence imaging of γH2AX and ACTA2 on confluent NHLF treated as in Fig. 2b. (**k, l**) Quantification of Mean Fluorescence Intensity (MFI) from (i) and (j). 3 biological replicates per condition. (**m**) NHLF cells were treated as of Fig. 2b and western blotted for γH2AX, and total H2AX. The western blots are representative of 3 independent experiments. **P < 0.01,***P < 0.001, ****P < 0.0001 were determined using a one-way ANOVA test with multiple comparisons by Tukey post-hoc analysis using GraphPad Prism 10.6 software.

### RAD51 Deficiency Suppresses mTOR Signaling

While our findings suggested that RAD51-mediated activity could be related to fibroblasts, the activation of fibroblasts and their subsequent resistance to apoptosis in pulmonary fibrosis are predominantly mediated through TGFβ signaling pathways, specifically via SMAD and non-SMAD mTOR pathways.^23–26^ Therapeutic interventions designed to inhibit these pathways aim to reduce fibroblast activation and restore apoptotic mechanisms to ameliorate lung fibrosis. To determine whether RAD51 can modulate TGFβ/SMAD/mTOR signaling and affect fibroblast behavior, NHLF cells were initially treated with siRNA targeting SMAD2, SMAD3, or a non-targeting control (**Fig. 3a**). Cells were then treated with inhibitors targeting MAP kinase (MEK), PI3K, AKT, and mTORC1 namely U0126, LY294002, MK2206, and rapamycin, respectively (**Fig. 3b**) and the induction of RAD51 by TGFβ was assessed. Although the phosphorylation of SMAD2 and SMAD3 is not dependent on RAD51 (**Fig. 3c, Fig. S4a and b**), the knockdown of SMAD2 or SMAD3, as well as the inhibition of MEK, PI3K, Akt, and mTORC1, abrogated the induction of RAD51 by TGFβ (**Fig. 3a and b**). This finding suggests that RAD51 expression is regulated by both canonical and non-canonical TGFβ signaling pathways. We investigated whether the loss of RAD51 led to a decrease in the phosphorylation of mTOR targets. The inhibition or silencing of RAD51 resulted in significantly lower levels of phosphorylated S6K (pS6K) and phosphorylated 4E-BP1 (p4E-BP1), indicating that RAD51 deficiency suppresses the mTORC1 pathway in fibroblasts (**Fig. 3d and e, Fig. S4c and d**).

**Fig. 3:**
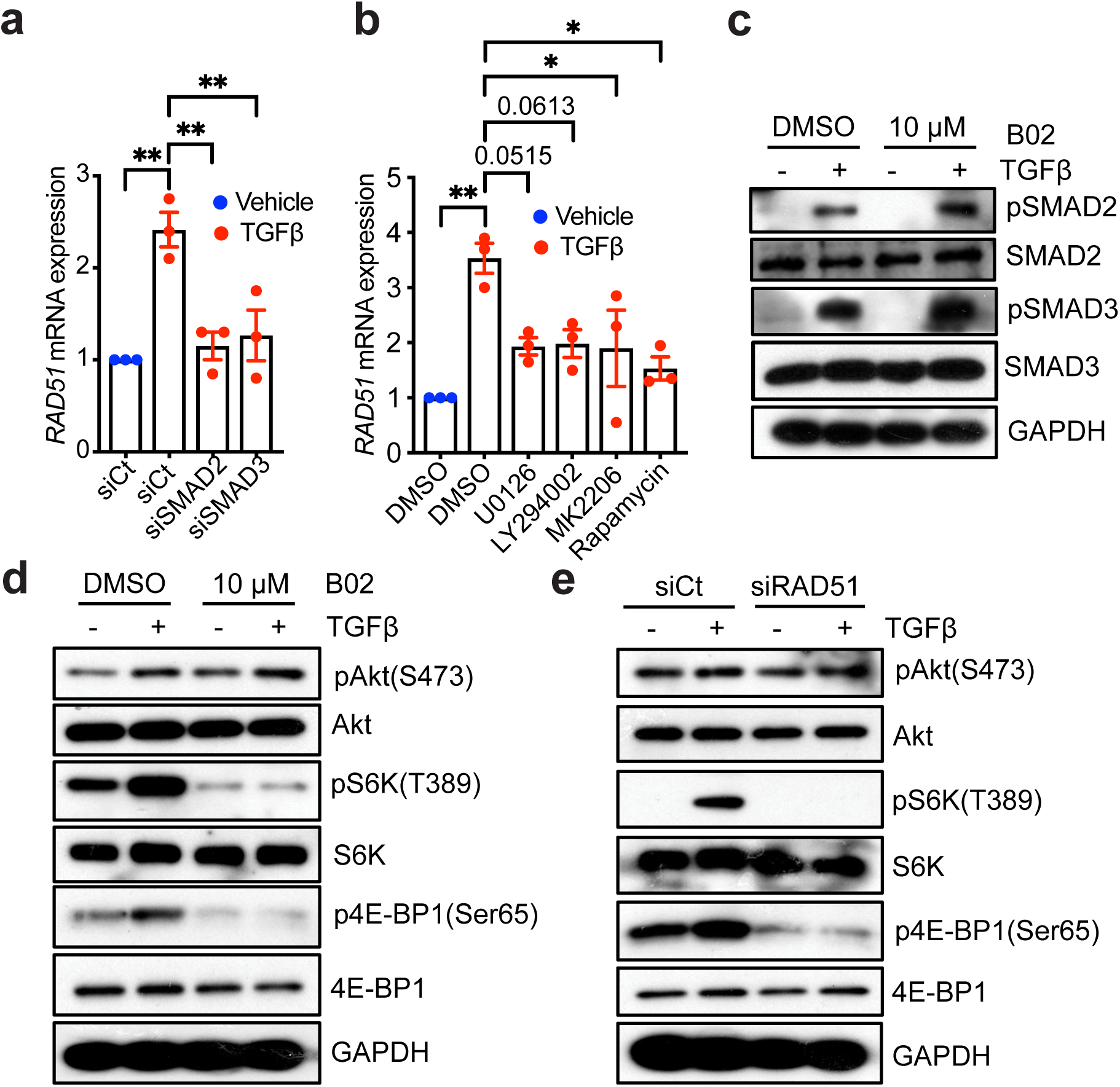
RAD51 Deficiency Suppresses mTOR Signaling. (**a**) NHLF cells with either a non-targeting control (siCt) or siRNA targeting SMAD2 or SMAD3, were treated with TGF-β (+; 5 ng/ml) or a vehicle (-). Subsequently, qPCR analysis of RAD51 was conducted 24 h after the treatment. (**b**) qPCR analysis for RAD51 post 24 h treatment with either the vehicle (-) or TGF-β (+) in the presence of 0.1% DMSO; MEK-ERK1/2 inhibitor U0126 (3 μM); PI3K inhibitor LY294002 (20 μM); Akt inhibitor MK2206 (300 nM); and mTORC1 inhibitor Rapamycin (100 nM). (**c**) NHLF cells were treated with B02 (10 μM) in the absence (−) or presence (+) of TGFβ for 6 h and subjected to western blot analysis for the indicated protein. (**d**) NHLF cells were treated with B02 (10 μM) and stimulated either in the absence (-) or presence (+) of TGFβ (5 ng/ml), and the indicated proteins were evaluated at 6 h. (**e**) NHLF cells were transfected with either scrambled control or RAD51 siRNA, followed by western blot analysis for the indicative proteins 6 h post treatment with TGF-β or a vehicle. All Western blots are representative of 3 separate experiments. Data presented in (a, b) indicate mean ± SEM with n = 3 independent experiments. *P < 0.05, **P < 0.01 were determined using one-way ANOVA followed by Tukey post-hoc test, using GraphPad Prism 10.6 software.

### RAD51 depletion disrupts mitochondrial metabolism

As inhibition of mTORC1 reduces ATP levels by slowing mitochondrial function and protein synthesis,^27^ we investigated whether loss of RAD51 diminishes ATP levels and affects the metabolic activity of fibrotic fibroblasts. We observed that depletion of RAD51 indeed led to reduced ATP levels in TGFβ activated fibroblasts (**Fig. 4a**). Given that ATP is predominantly produced via oxidative phosphorylation (OXPHOS) or glycolysis, we profiled the metabolic activity of these cells to determine whether RAD51 depletion affected either process using a real-time metabolite analyzer to measure the oxygen consumption rate (OCR) and extracellular acidification rate (ECAR) respectively. Inhibition of RAD51 with B02 resulted in decreased OCR under basal conditions, as well as upon treatment with oligomycin (an ATP synthase inhibitor), FCCP (an uncoupler of mitochondrial oxidative phosphorylation), and rotenone and antimycin A (inhibitors of mitochondrial electron transport complexes I and III, respectively), indicating impaired mitochondrial respiration (**Fig. 4b and c**). RAD51 inhibition significantly decreased HIF1α proteins supporting the positive impact of RAD51 on HIF1α protein regulation (**Fig. 4d**). As HIF1α is a master regulator of growth factor-dependent regulation of glucose metabolism,^28^ we accessed the influence of RAD51 on glycolysis. Inhibition of RAD51 inhibited glycolysis as reflected by suppressed ECAR (**Fig. 4e and f**). These observations demonstrate that RAD51 depletion reprograms cellular metabolism leading to reduced ATP production, OCR and ECAR. To further understand the impact of RAD51 depletion on mitochondrial respiration and glycolysis, we measured intracellular levels of glutamate and lactate. We found a significant reduction in glutamate levels in cells subjected to RAD51 inhibition (**Fig. 4g**). Although TGFβ significantly enhanced lactate production, we detected a reduction in lactate levels following RAD51 inhibition (**Fig. 4h**). Additionally, RAD51 inhibition led to decreased levels of NADP^+^/NADPH in TGFβ treated cells (**Fig. 4i**). This decrease in the NADP^+^/NADPH ratio indicates that the cell has a significant ability to produce NADPH, commonly to support reductive biosynthesis and combat elevated oxidative stress. RAD51 deficiency resulted in downregulation of GLS1 (glutaminase 1), which converts glutamine to glutamate in the mitochondria, suggesting a role of RAD51 in mitochondrial glutamine metabolism (**Fig. 4j and k, Fig. S5a and b**). These results demonstrate that RAD51 is both necessary and sufficient for the reprogramming of mitochondrial metabolism in fibrotic fibroblasts.

**Fig. 4:**
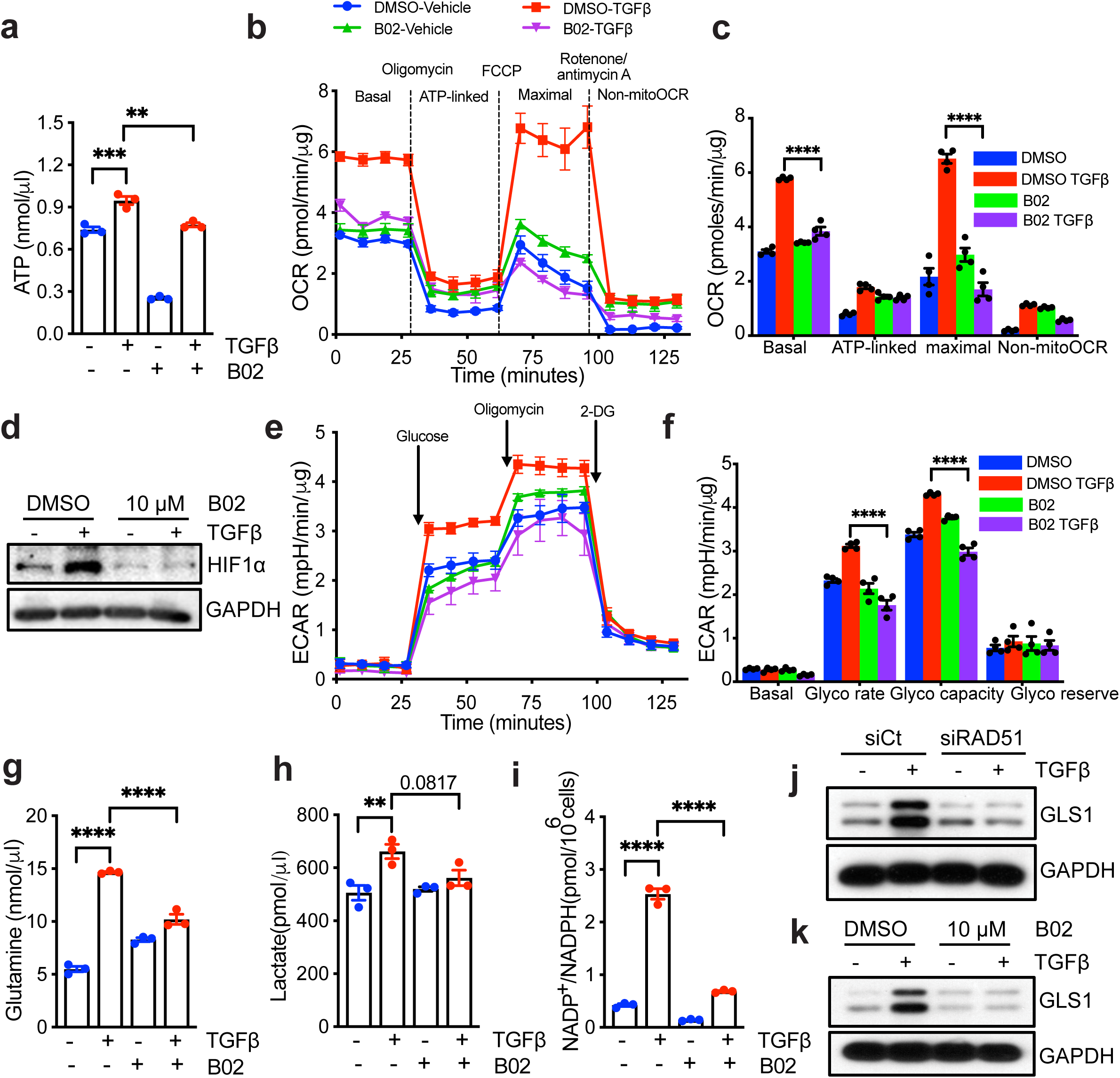
RAD51 depletion disrupts mitochondrial metabolism. (**a**) NHLF cells were treated with DMSO or B02. Following 48 h incubation (+/- 5 ng/ml TGFβ), lysates were prepared and cellular ATP level were measured. **(b, c)** Quiescent NHLF cells were treated for 24 h with either vehicle (DMSO) or B02 (10 μM), with or without TGFβ, and the oxygen consumption rate (OCR) was evaluated using the Seahorse XFp Cell Mito Stress Test Kit on a Seahorse XFp extracellular flux analyzer, with data presented as individual time points (b) and averages (c). (**d**) NHLF cells were treated as of Fig. 4a, lysates were prepared and western blotted for HIF1α. (**e, f**) NHLF cells were treated as described in 4b, and subsequent glycolytic activity was assessed through ECAR using the Seahorse XF Glycolysis Stress Test Kit, with results presented as individual time points (e) and averages (f) (**g**) Glutamine uptake assay in TGFβ-treated (48 h) NHLF cells demonstrates that TGFβ enhances glutamine uptake, which is subsequently inhibited by B02 (10 µM). (**h**) Quiescent NHLF cells were treated for 48 h to either DMSO or B02, with or without TGFβ, and intracellular lactate levels were measured across three independent experiments (n = 3). (**i**) NHLF cells were treated as described in Fig. 2b, and the levels of NADP^+^/NADPH were measured. (**j, k**) NHLF cells were treated as of Fig. 2a, b and western blotted for indicated protein. All data reflect the means ± SEM. Differences between groups were evaluated by one-way (a, g, h, i) or two-way (c, f) ANOVA with Tukey post-hoc analysis. **P < 0.01, ***P < 0.001, ****P < 0.0001.

### RAD51 inhibition induces fibroblast apoptosis

Pathologic IPF fibroblasts are resistant to apoptosis leading to enhanced survival ^10,29–32^ and mitochondria play a pivotal role in the resistance to apoptosis through energy metabolism.^33^ Considering that RAD51 depletion disrupted mitochondrial respiration with reduced ATP levels, we hypothesized that RAD51-deficient cells would exhibit increased sensitivity to apoptosis due to impaired energy metabolism. To test this possibility, NHLF were treated with B02 for 48 h and performed immunostaining or western blot analysis for cleaved caspase-3 to detect fibroblasts that are undergoing apoptosis. Compared to the control group, B02 treatment led to increased expression of cleaved caspase-3 in ACTA2^+^ fibroblasts (**Fig. 5a and b**). Additionally, primary human lung fibroblasts exhibited an increase in cleaved caspase 3 level (**Fig. 5c, Fig. S6a**) or caspase-3 activity following B02 treatment (**Fig. 5d**). In fibroblasts, the equilibrium between pro-apoptotic (BAX and BAK) and anti-apoptotic (BCL-2 and BCL-XL) factors governs apoptotic regulation.^10,34^ We hypothesized that blocking RAD51 activity would upregulate BAX and BAK and downregulate BCL-2 and BCL-XL, increase mitochondrial permeability transition pore (MPTP) opening and accelerate the release of mitochondrial cytochrome c (cyto c). This increased release of cyto c from mitochondria would enhance the activation of the caspase-3, thereby promoting apoptosis in myofibroblasts (**Fig. 5e**). We first examined whether RAD51 inhibition affected pro-apoptotic BCL-2 family members and isolated cytosolic and mitochondrial fractions from NHLF cells treated with B02. Inhibition of RAD51 increased the expression of pro-apoptotic factors BAK, BAX, PUMA, and BAD (**Fig. 5f, Fig. S6b**) in the mitochondrial fraction. We also observed that RAD51 inhibition increased acetylation of p53 at lysine 120 (K120) in mitochondria, a key modulator of cell fate decisions between apoptosis and cell cycle arrest (**Fig. 5g-i, Fig. S6c**).^35^ This posttranslational modification specifically triggers apoptosis by enhancing the binding affinity of p53 to pro-apoptotic gene promoters, such as PUMA.^36^ To test whether the upregulation of proapoptotic factors by RAD51 inhibition affects cyto c release from mitochondria, B02-treated cells exhibited higher cyto c content in the cytosolic fraction compared to the mitochondrial fraction (**Fig. 5j and k, Fig. S6d and e**), suggesting that RAD51 inhibition promotes cyto c release from mitochondria. Since MPTP opening is implicated in the release of cyto c and impeded in IPF fibroblasts conferring resistance to apoptosis,^37^ we investigated the role of RAD51 on fibroblast MPTP opening in control and TGFβ activated fibroblasts under vehicle or B02 treated conditions, using the CoCl_2_-calcein fluorescence-quenching assay by confocal microscopy as described.^37^ Calcein fluorescence revealed comparable levels in normal and TGFβ-activated fibroblasts without CoCl_2_, demonstrating comparable calcein incorporation (**Fig. S6f**). However, CoCl_2_ treatment resulted in reduced mitochondrial calcein fluorescence in B02-treated fibroblasts, indicating an increase in MPTP opening in these cells (**Fig. 5l and m**). These findings demonstrate that RAD51 inhibition activates the p53 pathway, amplifying its pro-apoptotic effects, leading to MPTP opening and cytochrome c release, which subsequently enhances caspase 3-mediated cell death in fibroblasts.

**Fig. 5:**
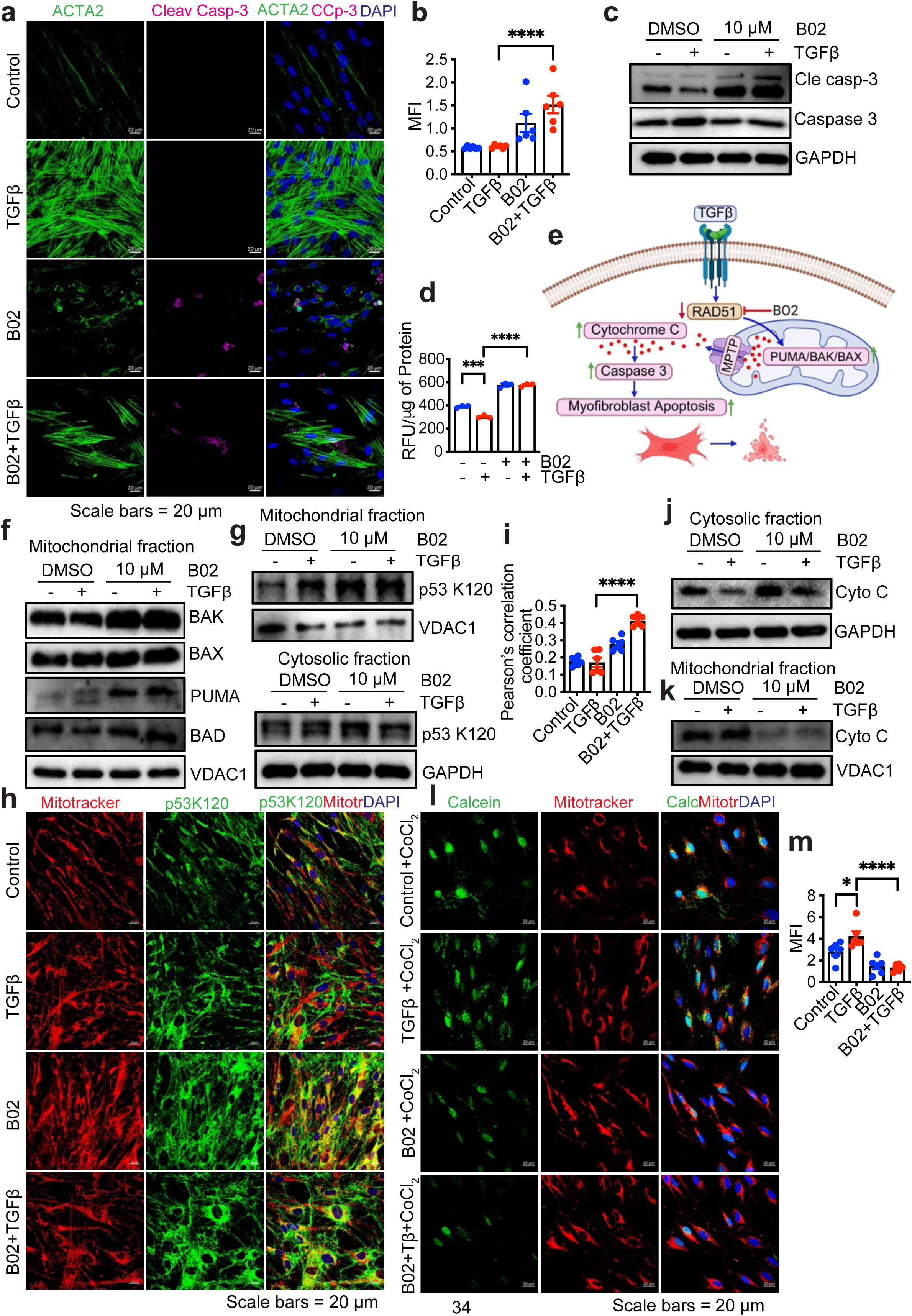
RAD51 inhibition induces fibroblast apoptosis. **(a)** Immunofluorescence staining of cleaved caspase-3 (pink), ACTA2 (green) and DAPI (blue) on confluent NHLF treated as in Fig. 2b. (**b**) Quantitative analysis of immunofluorescence of cleaved caspase 3. (**c**) Western blot analysis for cleaved caspase 3 and total caspase 3 protein in TGFβ treated (48 h) NHLF cells as in Fig. 2b. (**d**) NHLF cells were treated with B02 (10 μM) in the absence (-) or presence (+) of TGFβ (5 ng/ml), and apoptosis was measured by caspase-3 activity. Data reflect mean ± SEM of n = 3. (**e**) Schematic representation on the role of RAD51 in apoptotic signaling pathways that contribute to fibrosis. (**f**) NHLF cells were treated with B02 (10 μM) following 48 h incubation (+/- 5 ng/ml TGF-β), mitochondrial fractions were isolated and western blotted for indicated proteins. (**g, h**) Representative immunoblot of p53K120 in the mitochondrial or cytosolic fractions (g), or immunofluorescence staining of p53K120 in NHLF cells treated as described in Fig. 2b. (**i**) Quantification of p53K120 colocalization with mito tracker. At least five random fields were analyzed through ImageJ program. (**J, k**) Western blot of cytochrome c in cytosolic fractions (j) or mitochondrial fractions (k) of NHLF cells treated as of Fig. 2b. (**l**) Confocal microscopy of calcein and mito tracker on TGFβ activated NHLF +/- B02 following incubation with calcein-AM (1μM) in the presence of CoCl_2_ (1 mM). (**m**) Quantitative analysis of calcein immunofluorescence. Western blots are representative of 3 independent experiments. *P < 0.05, ***P < 0.001, ****P < 0.0001 were calculated by one-way ANOVA test with Tukey post-hoc test using GraphPad Prism 10.6 software.

### B02 reduces expression of fibrotic markers in Precision-cut lung slice (PCLS) model

We next sought to test the effect of RAD51 inhibition on fibroblast activity in PCLS, an organotypic model that retains the cellular heterogeneity and structural architecture of lung tissue.^38^ We first tested fibroproliferative responses ex vivo in wild type mice (mPCLS) or normal human (hPCLS) tissues (**Fig. 6a**). We cultured those PCLS for 4 days in the presence or absence of B02 and a fibrosis cocktail (TGFβ+TNFα). PCLS remained viable after 4 days of B02 treatment as measured by WST-1 proliferation assay (**Fig. S7a and b**). Immunofluorescence analysis demonstrated that TGFβ+TNFα treatment activated fibroblasts as shown by COL1 and ACTA2 expression while B02 suppressed its activation (**Fig. 6b and d, Fig. S7e and f**). We additionally prepared PCLS from the lungs of mice subjected to bleomycin challenge and evaluated the impact of B02 on the expression of fibrotic markers. The expression of profibrotic markers were elevated in the PCLS derived from bleomycin-challenged mice, whereas treatment with B02 inhibited this expression without compromising cell viability (**Fig. 6c and e, Fig. S7c**). PCLS treated with B02 showed reduced protein expression of fibrotic markers like FN, COL1 and ACTA2 (**Fig. 6f-h, Fig. S8a-c**). We prepared PCLS from explanted IPF lungs (**Fig. 6i**) and treated them with B02 for 72 h, followed by immunofluorescence and western blotting to assess profibrotic marker levels. The B02 treatment significantly reduced COL1 and other profibrotic expression compared to untreated lungs, indicating that RAD51 inhibition suppresses profibrotic protein expression in IPF lung (**Fig. 6j-l**) without impacting cell viability (**Fig. S7d**).

**Fig. 6:**
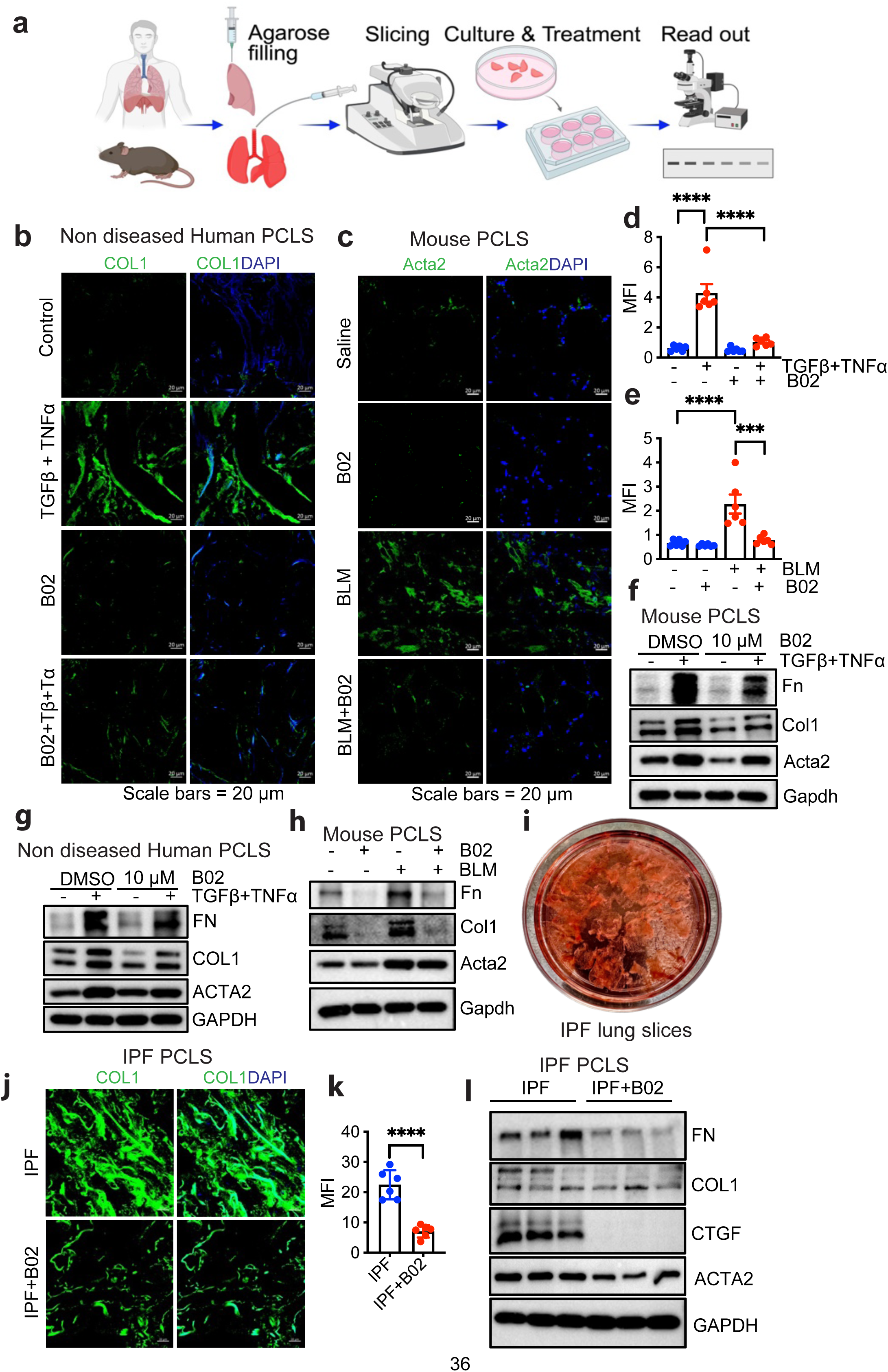
B02 reduces expression of fibrotic markers in Precision-cut lung slice (PCLS) model. **(a)** Schematic diagram of the workflow of PCLS. (**b**) Adult non-diseased human lung tissue samples were inflated, sliced into 250 µm thick sections using a vibrating microtome, then cultured in neurobasal media supplemented with 2% B27 and 1% P/S for 24 h, followed by treatment with 10 µM B02, 10 ng/ml TGFβ, and TNFα for an additional 72 h. After which immunofluorescence staining for the fibrotic marker COL1 was performed, with green indicating COL1 and blue indicating DAPI. (**c**) C57BL/6 mice were treated with either saline (Control) or bleomycin (BLM-2.5 U/Kg). On day 21, lungs were inflated, sectioned into 250 µm slices, and cultured in neurobasal media with 2% B27 and 1% P/S for 24 h. They were then treated with 10 µM B02 for 72 h and immunofluorescence staining was performed to detect the fibrotic marker Acta2 (green) and DAPI (blue). (**d, e**) Quantification of COL1 (d) and Acta2 (e) by ImageJ. (**f, g, h**) Western blot analysis of indicated protein from 8-12 week old C57BL/6 mice PCLS treated with or without B02 (10 μM) in the absence (−) or presence (+) of TGFβ+TNF-α (10 ng/ml each) for 72 h (f) or as mentioned on Fig. 6b (g) and Fig. 6c (h). (**i**) Representative image of ex-vivo culture of human 250 μM thick PCLS derived from IPF human lung. (**j-l**) Localization of COL1 was detected through immunofluorescence staining in lPF PCLS with antibodies for collagen 1 (green) and DAPI for nuclei (blue). Scale bars= 20 μm (j), immunofluoresce quatification of COL1 of j (k) or western blot analysis of profibrotic proteins (l) from precision cut IPF lung slices cultured with or without RAD51 inhibitor B02 for 72 h. Data are presented with n = 3 for mice, non-diseased human lung, or IPF lung. ***P < 0.001, ****P < 0.0001 were calculated by one-way ANOVA test with Tukey post-hoc test using GraphPad Prism 10.6 software.

### B02 treatment ameliorates bleomycin-induced fibrosis in a therapeutic mouse model

We extended our in vitro and ex vivo findings to a murine treatment model of lung fibrosis, where fibrosis was induced in 10-week-old C57BL/6 male and female mice by bleomycin (BLM) at a dose of 2.5 U/kg. Treatment was initiated on day 11, coinciding with the onset of fibrosis after the peak of inflammation. Based on previous studies showing that a B02 concentration of 50 mg/kg is optimal in tumor models, ^14,15^ mice received intraperitoneal injections of B02 (50 mg/kg) or 0.9% normal saline every two days, for a total of six injections, and were euthanized on day 23 (**Fig. 7a**). The level of peripheral blood oxygen saturation in mice breathing room air (SpO2), which serves as a non-invasive assessment of lung function and correlates with morbidity and mortality in the bleomycin-treated group, was measured. Lung compliance was also determined immediately post-euthanasia on day 23 using the flexiVent system. Both parameters indicated an improvement in lung function dependent on B02 treatment (**Fig. 7b and c**). These physiological findings were supported by enhanced H&E and trichrome staining results (**Fig. 7d and e, Fig. S9a**); reduced collagen accumulation, as evidenced by hydroxyproline levels (**Fig. 7f**); lower lung weight (**Fig. 7g**); and decreased expression of profibrotic markers (**Fig. 7h and i, Fig. S9b**). To further investigate the effects of B02 on fibroblast activation in bleomycin-induced lung tissue, we performed immunofluorescence staining for Cthrc1^+^ (Collagen triple helix repeat containing 1) pathologic fibroblasts.^39^ Compared to bleomycin-treated lungs, B02 treatment resulted in a lower number of Cthrc1^+^ fibroblasts (**Fig. 7j and k**). No evidence of systemic toxicity was observed in the liver, kidney, or blood parameters in B02-treated mice compared to control mice (**Fig. S10-12**). Taken together, these findings demonstrate that pharmacological inhibition of RAD51 using B02 during the fibrotic phase is sufficient to attenuate bleomycin-induced pulmonary fibrosis.

**Fig. 7:**
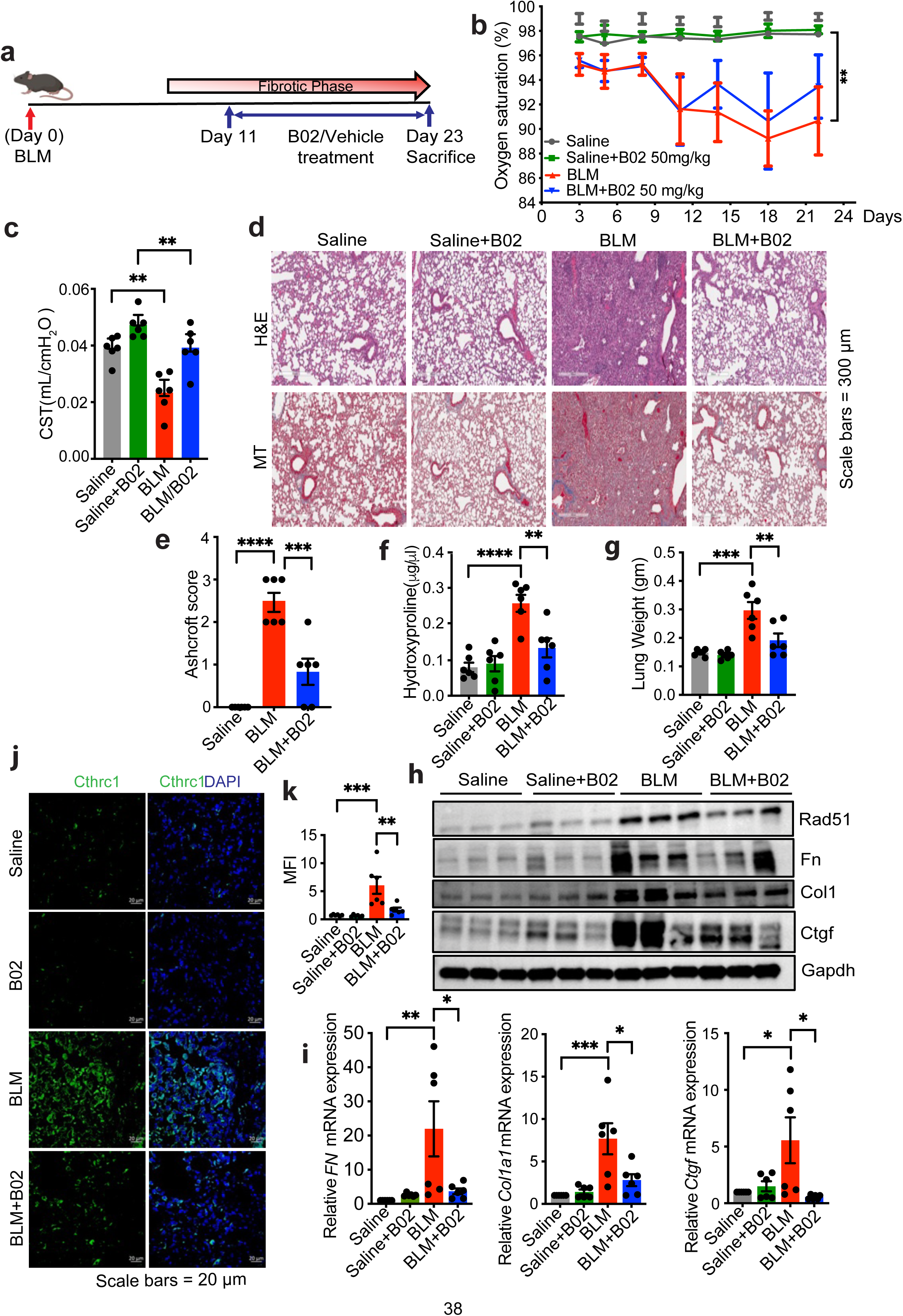
B02 treatment in a therapeutic murine model ameliorates bleomycin induced fibrosis. **(a)** Timeline for in vivo intratracheal administration of saline (Control) or bleomycin (BLM) at a dose of 2.5 U/Kg to C57BL/6 male and female mice. From day 11 to 21 mice were treated every two days, totaling six injections with either vehicle (PBS supplemented with 2% DMSO) or B02 (50 mg/Kg) by IP injection. On day 23, mice were euthanized. (**b**) Oxygen saturation levels were measured every third day following BLM administration. On day 23, mice were euthanized, and (**c**) lungs subjected to Flexivent determination. (**d, e**) Hematoxylin and Eosin (H&E) staining for histology and Masson’s trichrome (MT) for collagen deposition (blue) were performed (d) and the lung histopathology sections were blindly scored for the degree of fibrosis using Ashcroft method. Scoring was performed based on the following criteria: 0 (no fibrosis), 1 (focal/minimal fibrosis), 2 (multifocal/moderate fibrosis), 3 (confluent/severe fibrosis), Scale bars, 300 μm (e). (**f**) Total collagen levels were quantified using hydroxyproline assay. (**g**) Total lung weight. (**h**) Western blot analysis for fibrotic markers in murine lung tissue harvested on day 23. (**i**) qPCR analysis for fibrotic markers, *Fn, COL1a1 and Ctgf,* in murine lung tissue harvested on day 23. (**j**) immunofluorescence staining for Cthrc1, with green indicating Cthrc1 and blue indicating DAPI. (**k**) Quantification of Cthrc1 by ImageJ as mentioned in j. Data reflect means ±SEM of 6-7 mice for each group. Differences between groups were determined by two-way ANOVA test with Tukey post-hoc analysis (b) or one-way ANOVA with Tukey post-hoc analysis (c, e, f, g, i, k) using GraphPad Prism 10.6 software. *P < 0.05, **P < 0.01, ***P < 0.001, ****P < 0.0001.

## Discussion

DNA damage response pathways play an important role in determining the fate of fibroblasts when exposed to genotoxic stress by mediating either apoptosis or survival mechanisms.^11,12^ These pathways not only protect genomic integrity and facilitate DNA repair but also dictate cell fate decisions, thereby influencing the balance between cell survival and programmed cell death. An imbalance in these processes can contribute to pathological conditions.^40^ The expression of RAD51, a key protein involved in homologous recombination in DNA repair pathway, is upregulated in IPF fibroblasts and correlates with diminished capacity to undergo apoptosis in response to DNA-damaging agents.^11,12^ This adaptation allows fibroblasts to maintain their proliferative state in pathological fibrosis. Conversely, studies by Chen et al., have shown a significant decrease in RAD51 expression in alveolar epithelial cells following DNA damage induced by bleomycin, leading to impaired double-strand break repair and subsequent cellular senescence.^13^ These findings highlight the dual role of RAD51 in both DNA repair and cell survival under stress, which is particularly relevant in lung injury and fibrotic diseases where dysfunctional repair mechanisms exacerbate lung remodeling. Here, we extend this concept by (i) elucidating the relationship between RAD51 and the fibroproliferative activity of TGFβ, (ii) identifying its role in pathological myofibroblast differentiation, (iii) demonstrating how RAD51 inhibition reprograms fibroblast metabolism and promotes apoptosis, and (iv) exploring the therapeutic potential of targeting RAD51 to mitigate various aspects of fibrosis progression (**Fig. 8**).

**Fig. 8:**
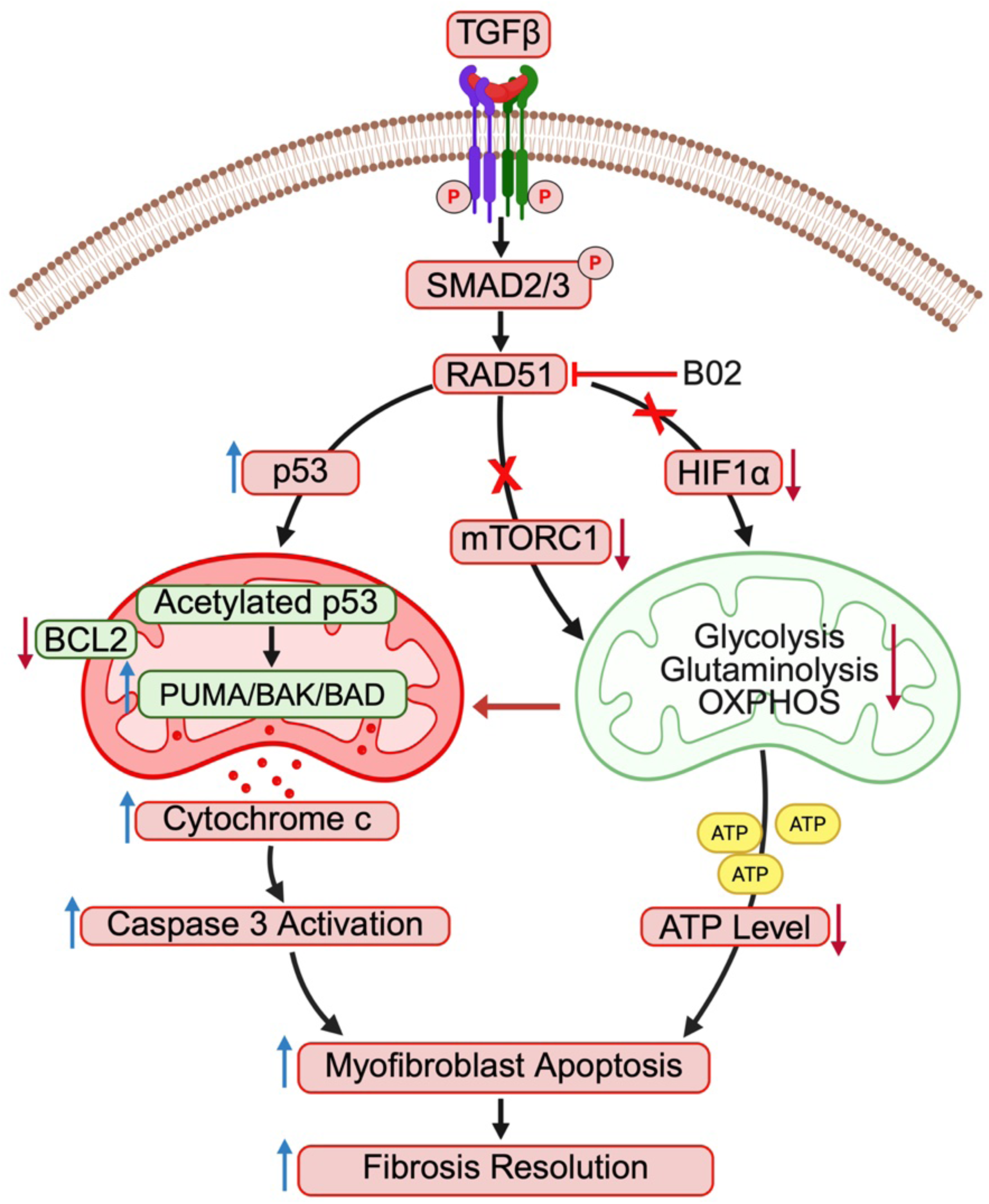
Schematic representation illustrating the role of RAD51 in the activation of fibroblasts. TGFβ-mediated upregulation of RAD51 decreases DNA damage, activates multiple pathways involved in fibroblast activation and drives metabolic reprogramming. RAD51 inhibition enhances p53 acetylation at lysine 120, increases mitochondrial expression of the proapoptotic proteins PUMA and BAK, and suppresses mTOR signaling and mitochondrial metabolism including glycolysis, glutaminolysis, and OXPHOS, thereby promoting myofibroblast apoptosis and fibrosis resolution.

Elevated RAD51 expression is associated with poor prognostic outcomes in various malignancies, as it promotes tumorigenesis and contributes to drug resistance.^41,42^ RAD51 is essential for repairing DNA damage induced by chemotherapeutic agents and ionizing radiation, enabling cancer cells to evade apoptosis and maintain genomic stability.^43^ Inhibition of RAD51 using the specific inhibitor B02 has been shown to increase the cytotoxicity of the chemotherapeutic drug cisplatin and enhance the therapeutic efficacy of receptor tyrosine kinase inhibitors in various cancer settings by preventing RAD51 from binding to single-stranded DNA and performing its DNA repair functions.^44,45^ Herein, we showed that RAD51 expression was upregulated in the lungs of IPF patients as well as in fibroblasts isolated from these patients. Blocking RAD51 activity using B02 or RAD51 expression using siRNA, inhibited TGFβ-mediated activation of fibroblasts and increased the expression of the DNA damage marker γ. This indicates that RAD51 is important for activating and protecting profibrotic fibroblasts through enhanced DNA repair. This observation aligns with previous research showing that in IPF fibroblasts, the transcription factor forkhead box M1 (FoxM1) upregulates RAD51 and BRCA2, thereby protecting cells from radiation-induced apoptosis and contributing to abnormal tissue repair mechanisms through increased DNA repair.^12^

Inhibition of RAD51 impacted various pathways associated with fibrotic lung diseases, highlighting its upstream role in many pathological processes. We determined that silencing or inhibiting RAD51 significantly decreased the phosphorylation of mTORC1 targets. Given that mTORC1 activation contributes to apoptosis resistance in IPF fibroblasts,^46^ and RAD51 silencing or inhibition enhances cell apoptosis in hepatocellular carcinoma,^47^ our results suggest that RAD51 inhibition leads to the accumulation of DNA damage in fibrotic fibroblasts which impairs mTORC1 activation, thereby increasing susceptibility to apoptosis.

DNA damage can induce metabolic reprogramming through the activation of the DNA damage response pathways. These pathways enhance nucleotide synthesis to facilitate DNA repair and increase cellular dependency on glucose and glutamine.^48^ Recent studies have highlighted the significance of metabolic reprogramming in activated fibroblasts, emphasizing their requirement for energy and biosynthetic precursors to support rapid proliferation and matrix production.^49–51^ Our research has demonstrated that RAD51 deficiency leads to metabolic reprogramming by impairing mitochondrial OXPHOS. As enhanced OXPHOS in TGFβ-stimulated human lung fibroblasts is essential for their survival and supports various fibrogenic responses by providing ATP, tricarboxylic acid cycle intermediates, and ROS,^52^ our findings underscore a significant role of RAD51 in mitochondrial metabolic reprogramming and glutamine metabolism. Additionally, our study supports the hypothesis that RAD51 facilitates myofibroblast glycolysis through the regulation of HIF1α, leading to increased glycolysis and apoptosis resistance. This suggests that RAD51 deficiency introduces metabolic vulnerabilities by reducing mitochondrial respiration, glycolysis, and glutaminolysis, which could potentially be therapeutically beneficial.

The accumulation of apoptosis-resistant, activated myofibroblasts within the pulmonary interstitium exacerbates fibrotic processes and impairs normal lung architecture and function. The equilibrium between pro-apoptotic and anti-apoptotic BCL-2 family proteins in fibroblasts determines their survival and susceptibility to apoptosis.^34^ Myofibroblasts have been shown to significantly downregulate pro-apoptotic proteins such as BAX and BAK, while upregulating anti-apoptotic proteins like BCL-2, thereby promoting fibrosis.^10,19^ Given that profibrotic fibroblasts resist apoptosis and rely on RAD51 for survival, we investigated the role of RAD51 in apoptosis. Mechanistically, our data demonstrate that inhibition of RAD51 results in the translocation of pro-apoptotic proteins (BAK, PUMA, BAD) from the cytosol to the mitochondria. This is mediated by enhanced acetylation of p53 at lysine 120, which subsequently activates BAK/PUMA, leading to the opening of the mitochondrial permeability transition pore, the release of cyto c, and caspase 3-mediated increased cell death in fibroblasts. K120 acetylation is a crucial post-translational modification that functions as a switch for p53, steering the cell towards apoptosis via the mitochondrial pathway instead of causing cell cycle arrest.^53^ This modification enhances p53’s antifibrotic activity. This is consistent with a previous study that demonstrated miR-106b-5p selectively eliminated senescent cells through p53-mediated upregulation of PUMA.^36^

We demonstrated that inhibition of RAD51 could alter lung fibrosis pathways in ex vivo and in vivo fibrosis models, suggesting RAD51 as a therapeutic target. PCLS serves as an ex vivo treatment model that maintains the three-dimensional intricate architecture and cellular composition of lung tissue to enable the cellular interactions associated with fibrotic processes. PCLS can serve as a predictive model for therapeutic efficacy in the lung, as illustrated by our evaluation of RAD51 inhibitor B02 in human and murine PCLS systems. Validation of the PCLS model for testing RAD51 were observed in a murine bleomycin model of pulmonary fibrosis. In our studies, B02 treatment had no detectable influence in normal tissue but RAD51 inhibition downregulated total collagen content and profibrotic markers expression and improved peripheral blood oxygen saturation on room air and lung compliance. B02 or related RAD51 inhibitors have not yet been used in clinical trials, though our data and results from preclinical cancer studies suggest a potential benefit.

There are limitations associated with this study. While it can be argued that RAD51-dependent repair is a universal double stranded break repair mechanism and its deletion or inhibition would be toxic to normal cells in gut, skin, mucosa and bone marrow, we observed no apparent toxicity in our short term in vitro and in vivo studies. Another potential limitation is that the inhibition of RAD51 by B02 in bleomycin injected mice is not exclusive to fibroblasts, and the impact of B02 on immune and other cells has not been determined. Future research will involve a mouse genetic model to knock out RAD51 globally compared to conditionally targeting fibroblasts to evaluate animal phenotypes and progression of the disease.

Together our findings defined a comprehensive set of models to integrate the proliferative and antiapoptotic functions of RAD51 in TGFβ-driven fibrosis. As single agent therapies for pulmonary fibrosis are highly ineffective^5^, we propose that perhaps a combination of DNA damage response pathway inhibition (using B02) with antiapoptotic blockade (using BCL-2 inhibitor navitoclax) could be a potentially effective therapy against fibrosis. This approach would provide compelling rationale to extend the concept to a Phase I clinical trial testing the premise that combination therapy (i.e., each at suboptimal concentrations) for improved efficacy with reduced toxicity. This is the first extensive study of the relationship between DNA repair protein RAD51 and mTORC1 signaling, mitochondrial metabolism and apoptosis in fibrosis. The efficacy of RAD51 inhibition in fibrosis suggests that targeting DNA repair is a viable approach in fibrosis treatment and provides a framework for paradigm-shifting therapies broadly targeting fibroblast function together with deranged fibroblast metabolism.

## Author contributions

M.C., and A.H.L., designed research; R.K.M., A.K.S., K.J.S., L.M., J.R.K., and M.C., performed the experiments. M.C., R.K.M., and A.K.S., wrote the manuscript. All authors analyzed data and edited the manuscript. R.K.M., and A.K.S., contributed equally to this work.

## Acknowledgements

We thank Dr. Carol Feghali-Bostwick (Medical University of South Carolina, Charleston, SC) for providing primary lung fibroblasts from healthy individuals and IPF patients. We also thank Dr. Steven Brody for critically reading the manuscript. This research was supported by grants from the “Boehringer Ingelheim Discovery Award in Idiopathic Pulmonary Fibrosis/Interstitial Lung Disease” (M.C.), the NIH/NHLBI (R56HL158549 and R01HL167732) (M.C.), ALA DA-1277949 (J.R.K) and NIH K08 HL159418 (J.R.K). Figures (Fig. 5e and Fig. 8) were created using BioRender. During the preparation of this manuscript, the authors used ChatGPT solely to improve the language and readability of the text.

## Funding

Boehringer Ingelheim Discovery Award in IPF/ILD (MC), NIH R56HL158549 (MC), NIH R01HL167732 (MC), ALA DA-1277949 (J.R.K) and NIH K08 HL159418 (J.R.K).

## Notes

### Competing Interest Statement

The authors have declared no competing interest.

