## Supplemental File for "Targeting the DNA damage repair protein RAD51 alters fibroblast metabolism and enhances apoptosis in pulmonary fibrosis"

**Supplementary Figure S1**

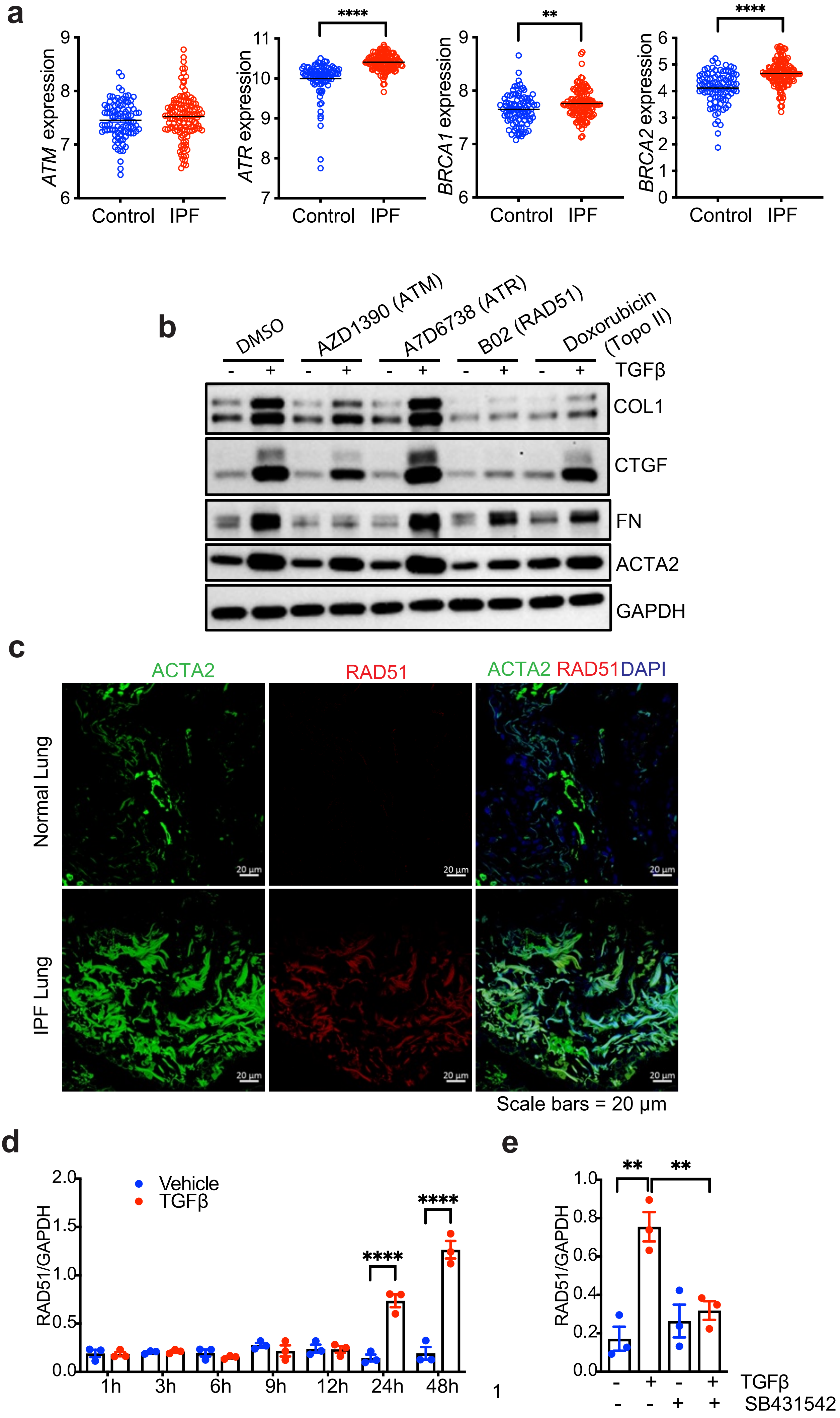

**Supplementary Figure S1. Upregulation of RAD51 in pulmonary fibrosis and quantitation of TGFβ induced RAD51 accumulation.** (**a**) *ATM, ATR, BRCA1, BRCA2* gene expression from 122 IPF and 91 controls patients of Lung Genomics Research Consortium (LGRC) microarray dataset (accession number GSE47460). (**b**) Quiescent NHLF cell lines were treated with either DMSO or ATM inhibitor (AZD1390), ATR inhibitor (A7D6738), RAD51 inhibitor (B02), Topoisomerase II inhibitor (Doxorubicin) in the absence (−) or presence (+) of TGFβ for 48 h and western blotted for the indicated proteins. GAPDH was used as a loading control. (**c**) Localization of RAD51 was identified using immunofluorescence staining in lung tissue sections obtained from both control and IPF patients, utilizing antibodies for ACTA2 (green), RAD51 (red), and DAPI for nuclei (blue). Co-localization can be seen in the merged images (far right side) for lung tissues from both non-IPF and IPF patients. Scale bars= 20 µm. **(d, e)** Ratios of RAD51 to GAPDH in NHLF cells were processed as depicted in Figures 1j and 1k. n = 3 independent experiments. Differences between groups were determined by two-way (d) or one-way (a, e) ANOVA test with Tukey post-hoc analysis using GraphPad Prism 10.6 software. **P < 0.01, ****P < 0.0001.

**Supplementary Figure S2**

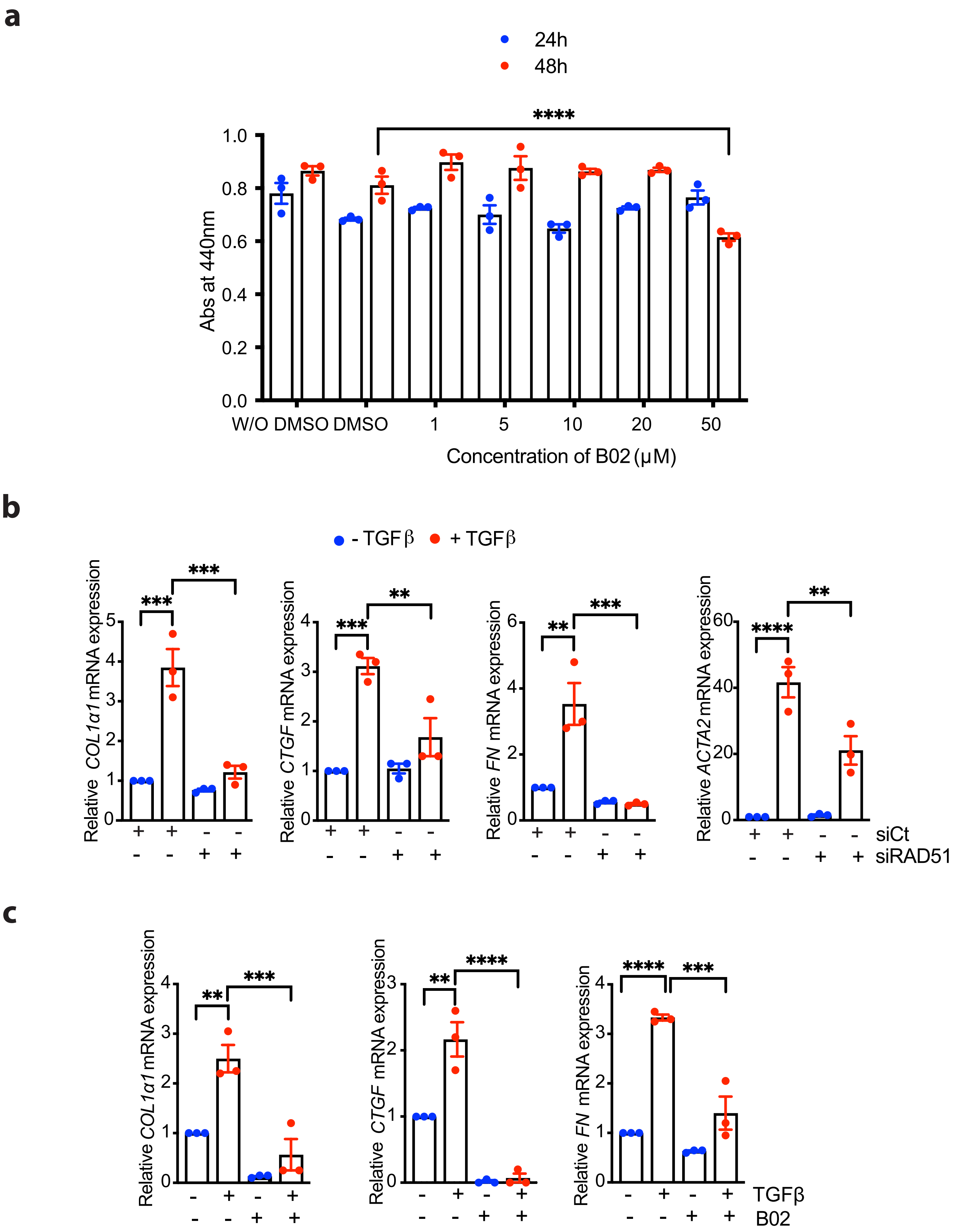

**Supplementary Figure S2. Profibrotic TGFβ signaling is dependent on RAD51.** (**a**) The WST-8 cell proliferation assay showed the maintained viability of NHLF cells following 24 h or 48 h incubation with RAD51 specific inhibitor B02 (1μM, 5μM, 10μM, 20μM and 50μM). 100μl/ml WST-8 reagent was added to each well containing NHLF and incubated for additional 2 h. Absorbance was measured at 440 nm using a microplate reader. Wells containing culture medium and WST-8 but without cells served as background controls. (**b**) NHLF cells were transfected with either a non-targeting control (siCt) or an siRNA targeting RAD51 (siRAD51). Following transfection, cells were treated with either a vehicle (-) or TGFβ (+) at a final concentration of 5 ng/ml. After 48 h of incubation, qPCR analyses were performed to measure the expression levels of indicated profibrotic molecules. (**c**) Quiescent NHLF cells were pretreated for 1 h with either 0.1% DMSO or 10 μM B02 (a RAD51-specific inhibitor) before the addition of vehicle or TGF-β (5 ng/ml). After a 48 h incubation period, qPCR analysis was performed to measure the expression levels of the specified profibrotic genes. n = 3 independent experiments. Differences between groups were determined by one-way ANOVA test with Tukey post-hoc analysis using GraphPad Prism 10.6 software. **$P < 0.01$, ***$P < 0.001$, ****$P < 0.0001$.

**Supplementary Figure S3**

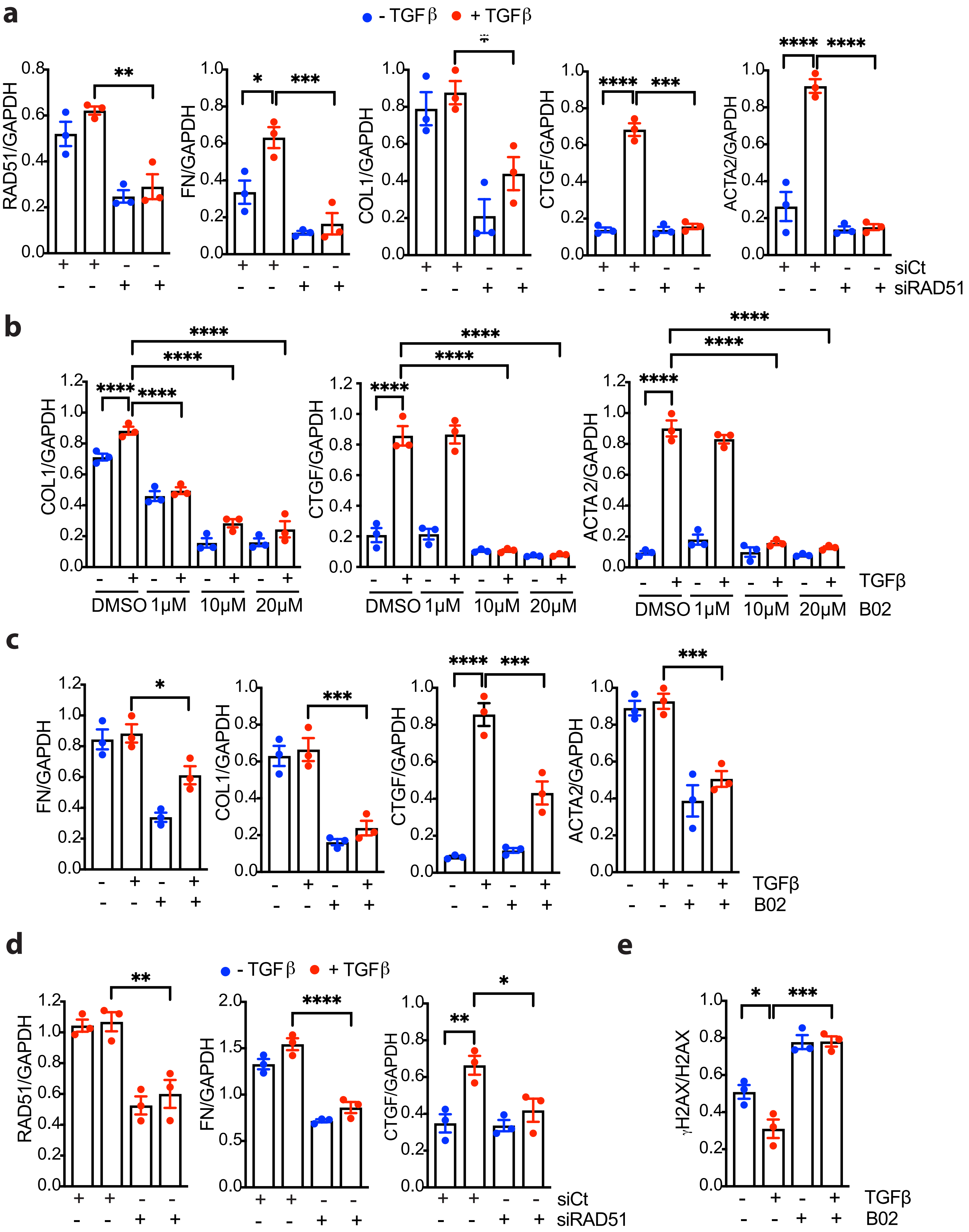

**Supplementary Figure S3. Quantification of RAD51 and profibrotic markers in NHLF or IPF fibroblasts following RAD51 knockdown or inhibition. (a-d)** Ratios of RAD51 and profibrotic proteins to GAPDH processed as in Fig. 2a, 2b, 2c and 2d. **(e)** Ratios of $\gamma$H2AX to H2AX processed as Fig. 2m. n = 3 independent experiments. All Data reflect the means ± SEM. Differences between groups were evaluated by one-way ANOVA with Tukey post-hoc analysis using GraphPad Prism 10.6 software. *P < 0.05, **P < 0.01, ***P < 0.001, ****P < 0.0001.

**Supplementary Figure S4**

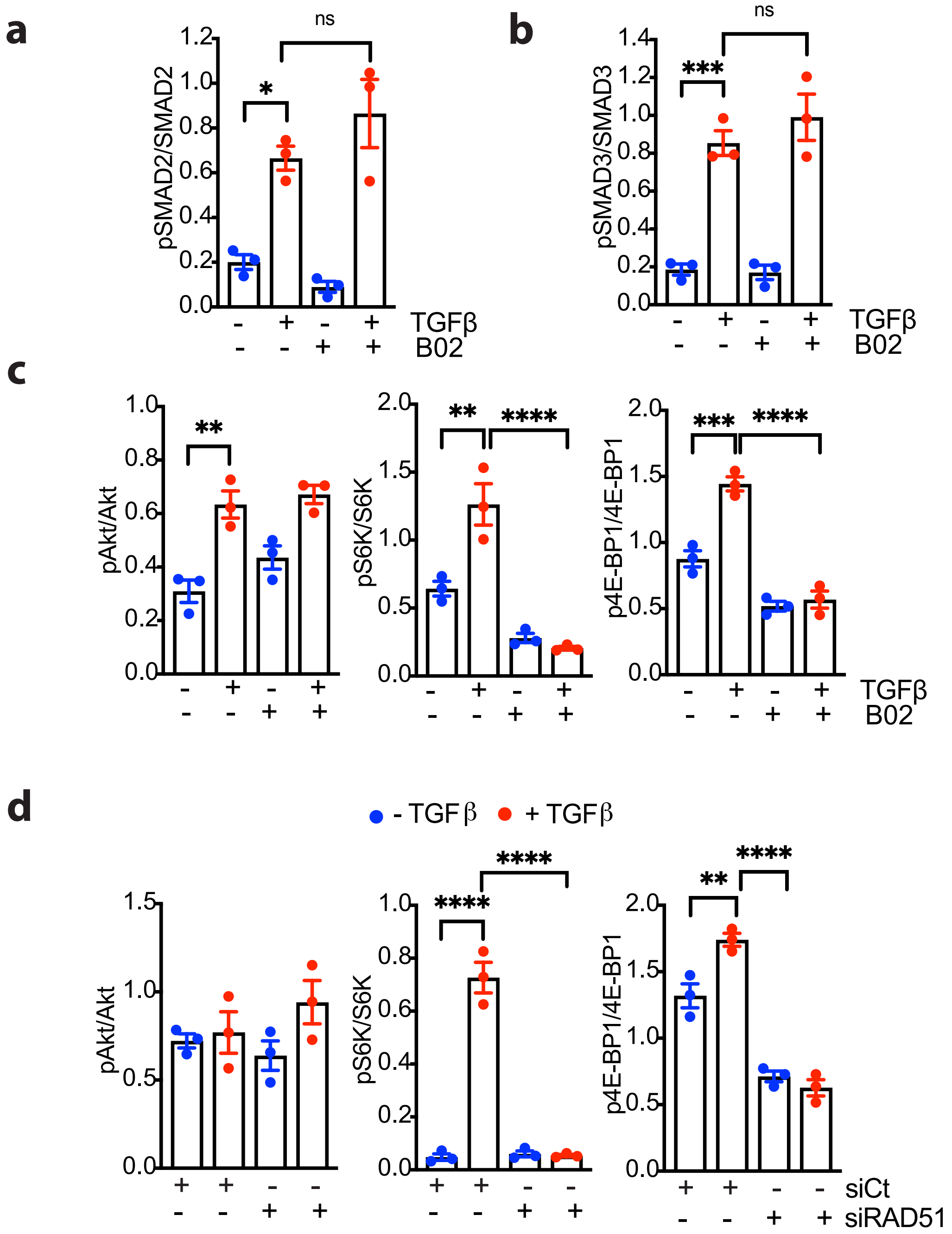

**Supplementary Figure S4. Role of RAD51 in SMAD and mTOR signaling. (a, b)** Ratios of pSMAD2 to SMAD2 (a) or pSMAD3 to SMAD3 (b) processed as in Fig. 3c. **(c, d)** Abundances of pS6K, pAkt and p4E-BP1 relative to total S6K, Akt or 4E-BP1, respectively as in Figure 3d and e. n = 3 independent experiments. *P < 0.05, **P < 0.01, ***P < 0.001, ****P < 0.0001.

**Supplementary Figure S5**

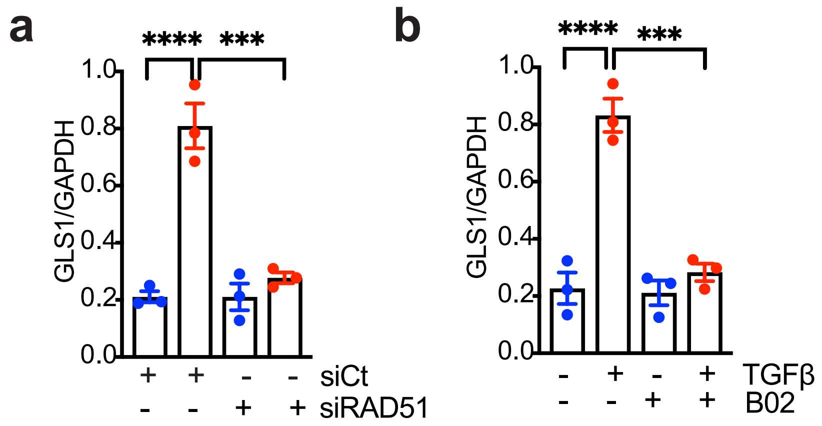

**Supplementary Figure S5. Quantification of metabolic target in NHLF. (a, b)** Quantification of GLS1 to GAPDH in NHLF cells processed as in Fig. 4j and k. n = 3 experiments. All Data reflect the means ± SEM. Differences between groups were evaluated by one-way ANOVA with Tukey post-hoc analysis. ***P < 0.001, ****P < 0.0001.

**Supplementary Figure S6**

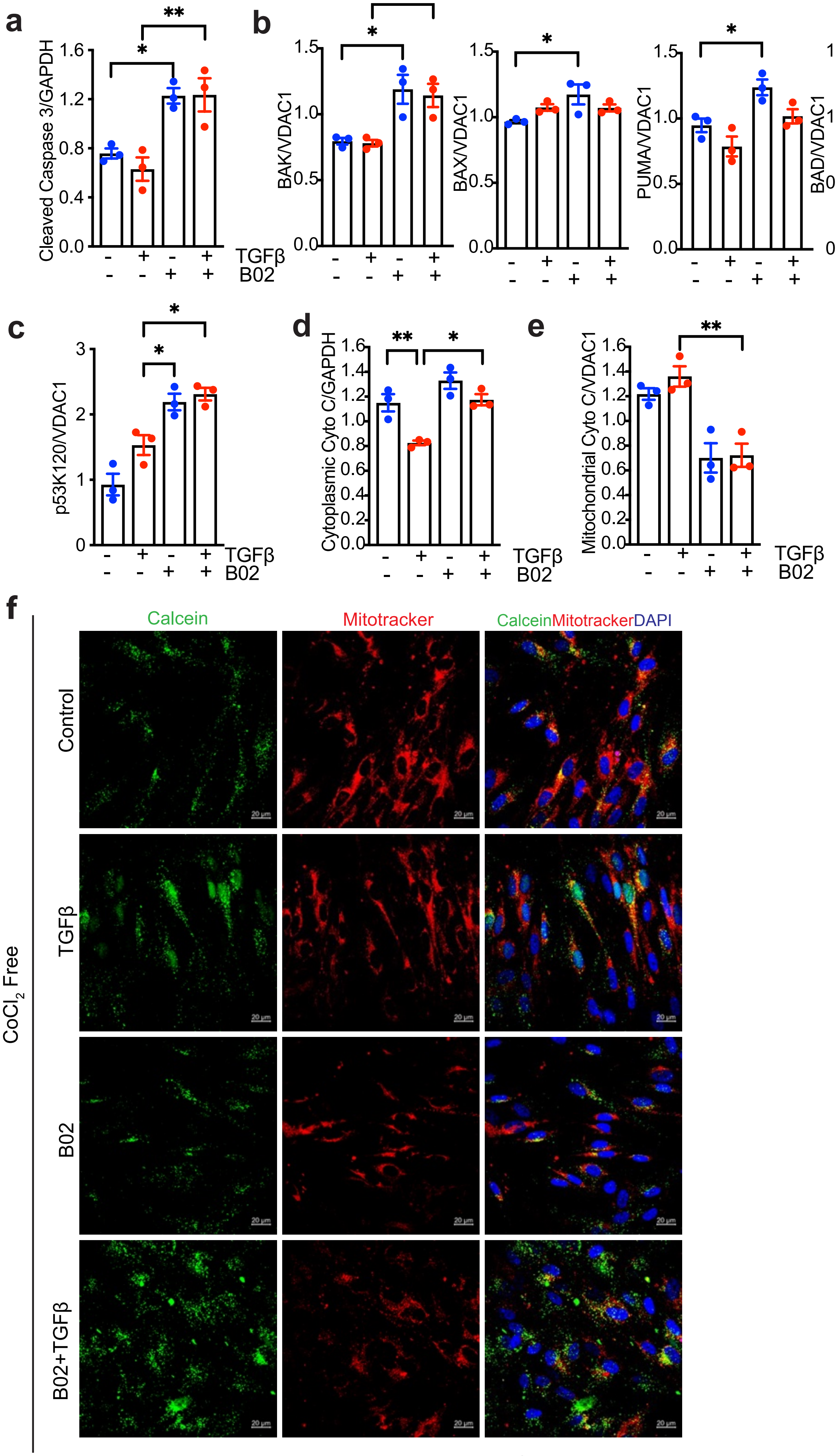


Scale bars = 20 μm

**Supplementary Figure S6. Quantitation of apoptosis markers in NHLF cells after RAD51 inhibition. (a-c).** Ratios of cleaved caspase 3 to GAPDH (a), proapoptotic factors to VDAC1 (b) and p53K120 to VDAC1 (c) as of Fig. 5c, f and g (mitochondrial fraction). (**d, e**) Cytochrome c to GAPDH in cytosolic fraction (d) and cytochrome c to VDAC1 in mitochondrial fraction (e), as mentioned in Fig. 5j and k. **(f)** Confocal microscopy of calcein and mitotracker was performed on TGFβ activated NHLF +/- B02 after incubation with calcein-AM (1$\mu$M) without the presence of $CoCl_2$. All Data reflect the means ± SEM. Differences between groups were evaluated by one-way ANOVA with Tukey post-hoc analysis. *$P < 0.05$, **$P < 0.01$.

**Supplementary Figure S7**

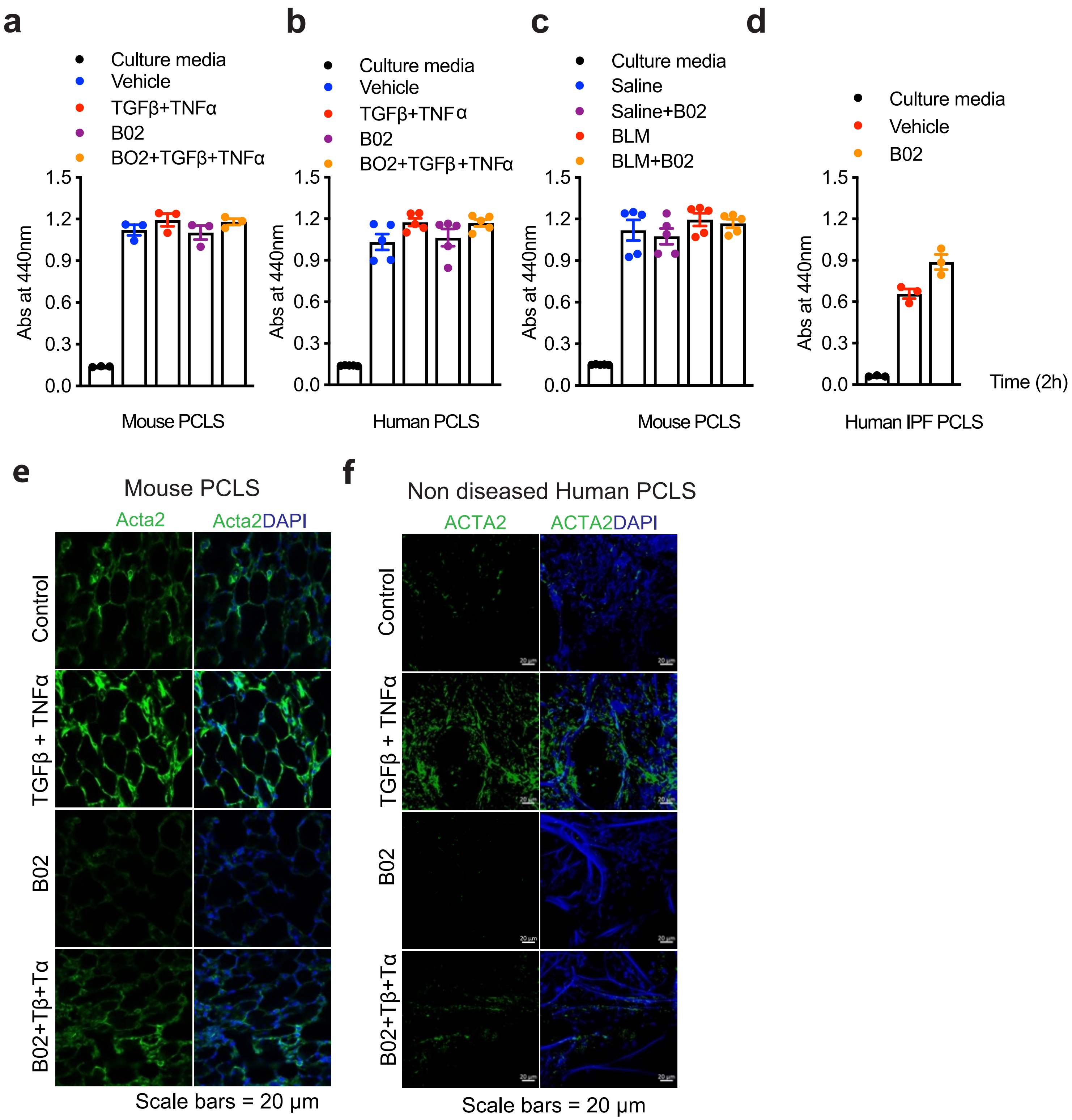

**Supplementary Figure S7. Quantification of viability and profibrotic markers in mouse and human PCLS treated with B02. (a-d)** Wst-8 assay showed the maintained viability of mouse PCLS (a, c) or human PCLS (b, d). Following 72 h incubation with indicated reagents, 100µl/ml WST-8 reagent was added to each well containing PCLS and incubated for 2 h. Absorbance was measured at 440 nm using a microplate reader. Wells containing culture medium and WST-8 but without slices served as background controls. **(e, f)** 8–12-week C57BL/6 mice lung (e) or adult non-diseased human lung tissue samples (f) were inflated and sliced into sections 250 µm thick using a vibrotome. These sections were then cultured in neurobasal media for 24 h, followed by treatment with 10 µM B02, 10 ng/ml TGFβ, and TNFα for an additional 72 h. Subsequently, immunofluorescence staining for the fibrotic marker ACTA2 was conducted, with green representing ACTA2 and blue representing DAPI. Data are representative of 3 independent experiments.

### Supplementary Figure S8

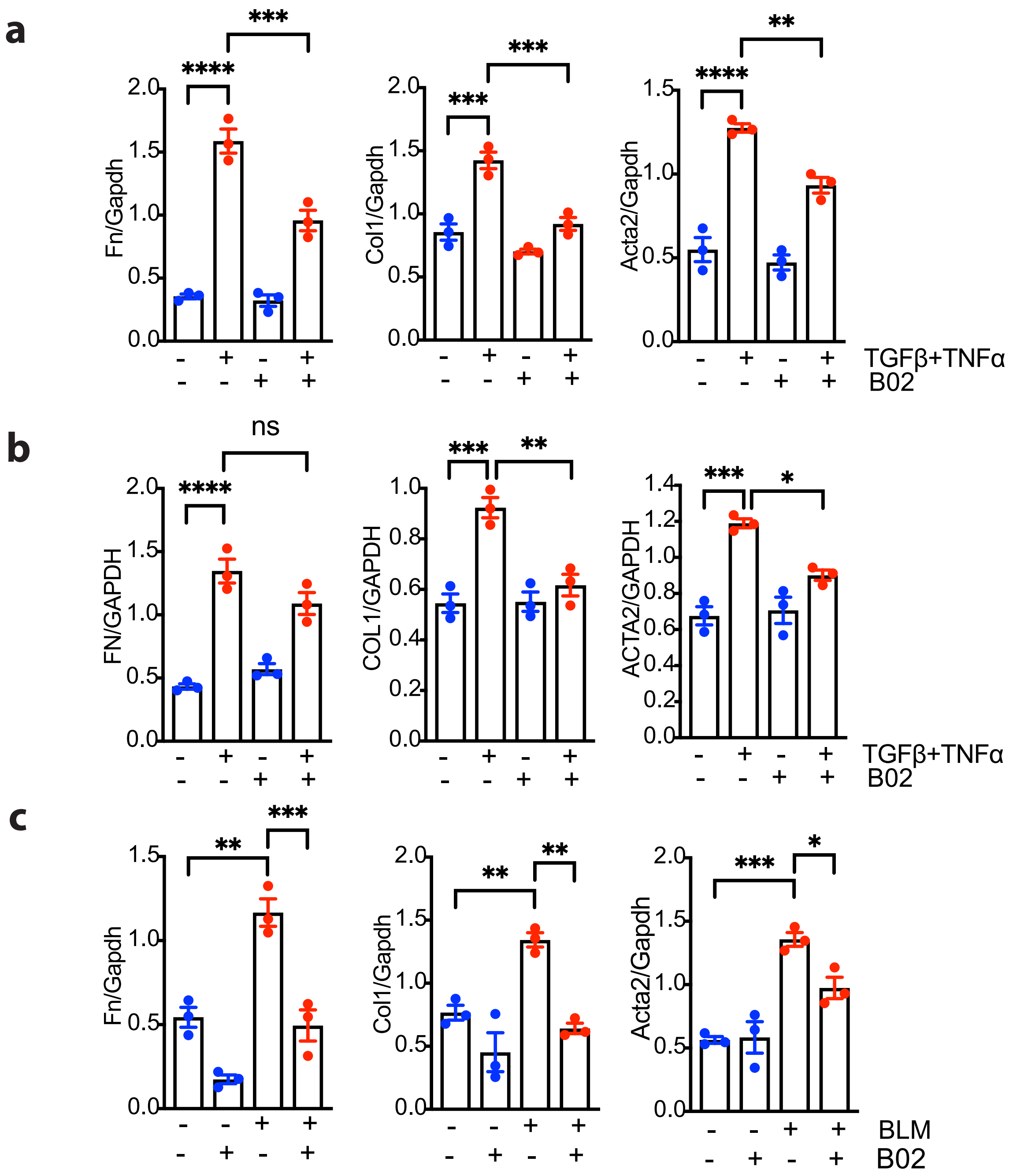

**Supplementary Figure S8. Quantitation of profibrotic markers in PCLS after RAD51 inhibition. (a, b, c)** Ratios of profibrotic marker (FN, Col1, ACTA2) to GAPDH processed as in Fig. 6f, g and h. n = 3 independent experiments. All Data reflect the means ± SEM. Differences between groups were evaluated by one-way ANOVA with Tukey post-hoc analysis. *P < 0.05, **P < 0.01, ***P < 0.001, ****P < 0.0001.

**Supplementary Figure S9**

**a**

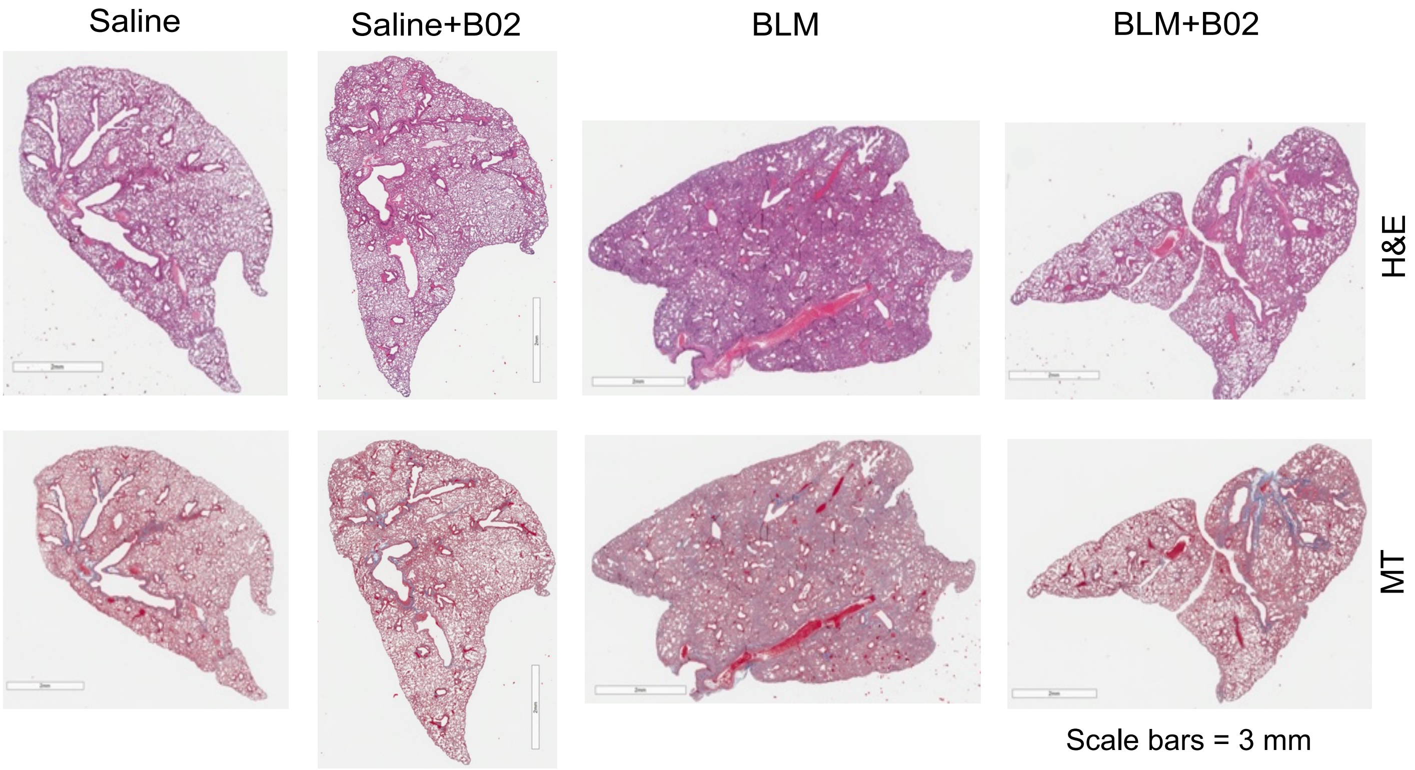


**b**

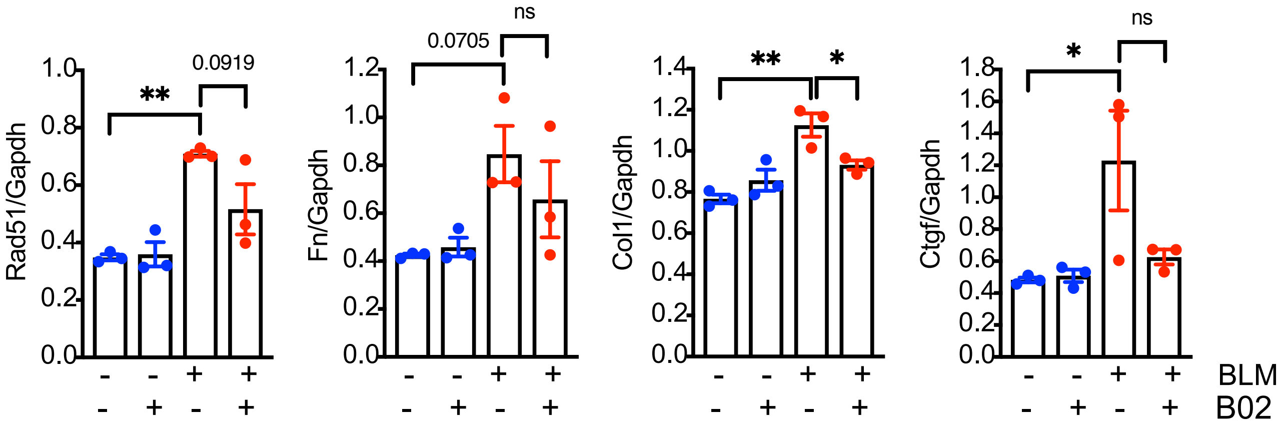

**Supplementary Figure S9. B02 attenuates bleomycin-induced pulmonary fibrosis affecting lung structure**. **(a)** Mice were treated as depicted in Fig. 7a, and Hematoxylin and Eosin (H&E) staining for histological analysis along with Masson's trichrome (MT) staining for collagen deposition (indicated in blue) were conducted following euthanasia on day 23. Representative images from 6-7 mice are presented. Scale bars, 3 mm. **(b)** Quantification of Rad51, Fn Col1, and Ctgf relative to Gapdh was processed as shown in Fig. 7h. The data are represented as mean ± SEM with n = 3. Differences among groups were assessed using one-way ANOVA with Tukey post-hoc analysis. *$P < 0.05$, **$P < 0.01$.

**Supplementary Figure S10**

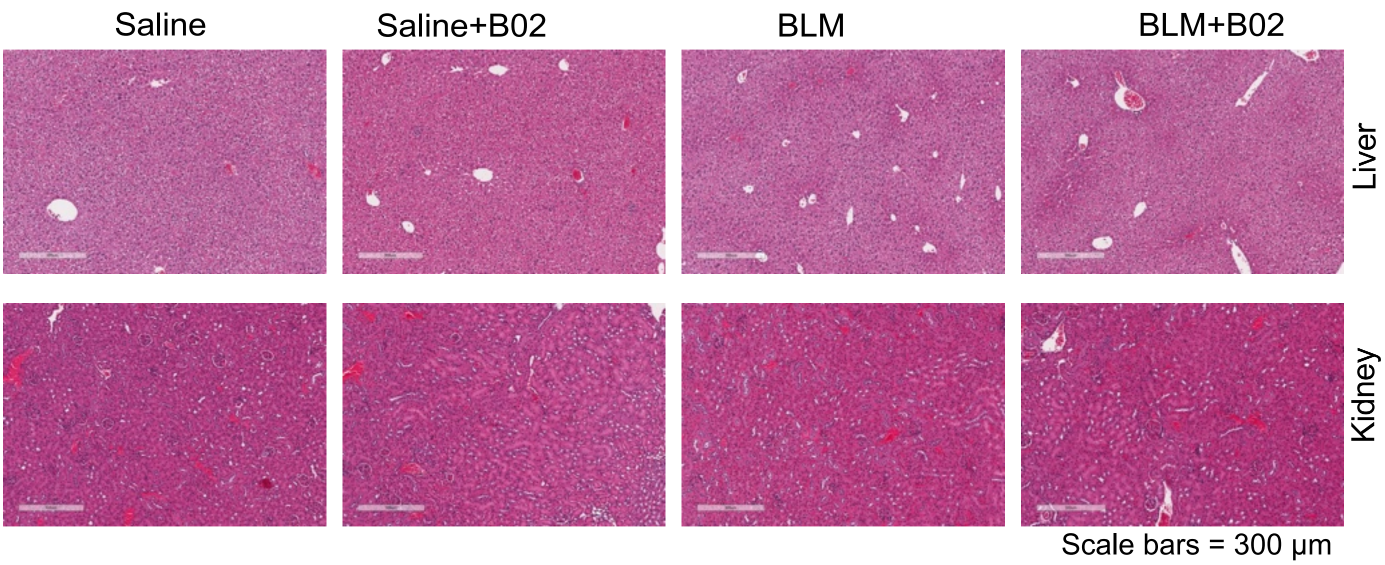

**Supplementary Figure S10. Pathology of the liver and kidneys was comparable in mice that were subjected to treatment with either B02 or the vehicle**. H&E staining was performed on the histology of the liver and kidney.  Data are presented as of n = 6 mice.

**Supplementary Figure S11**

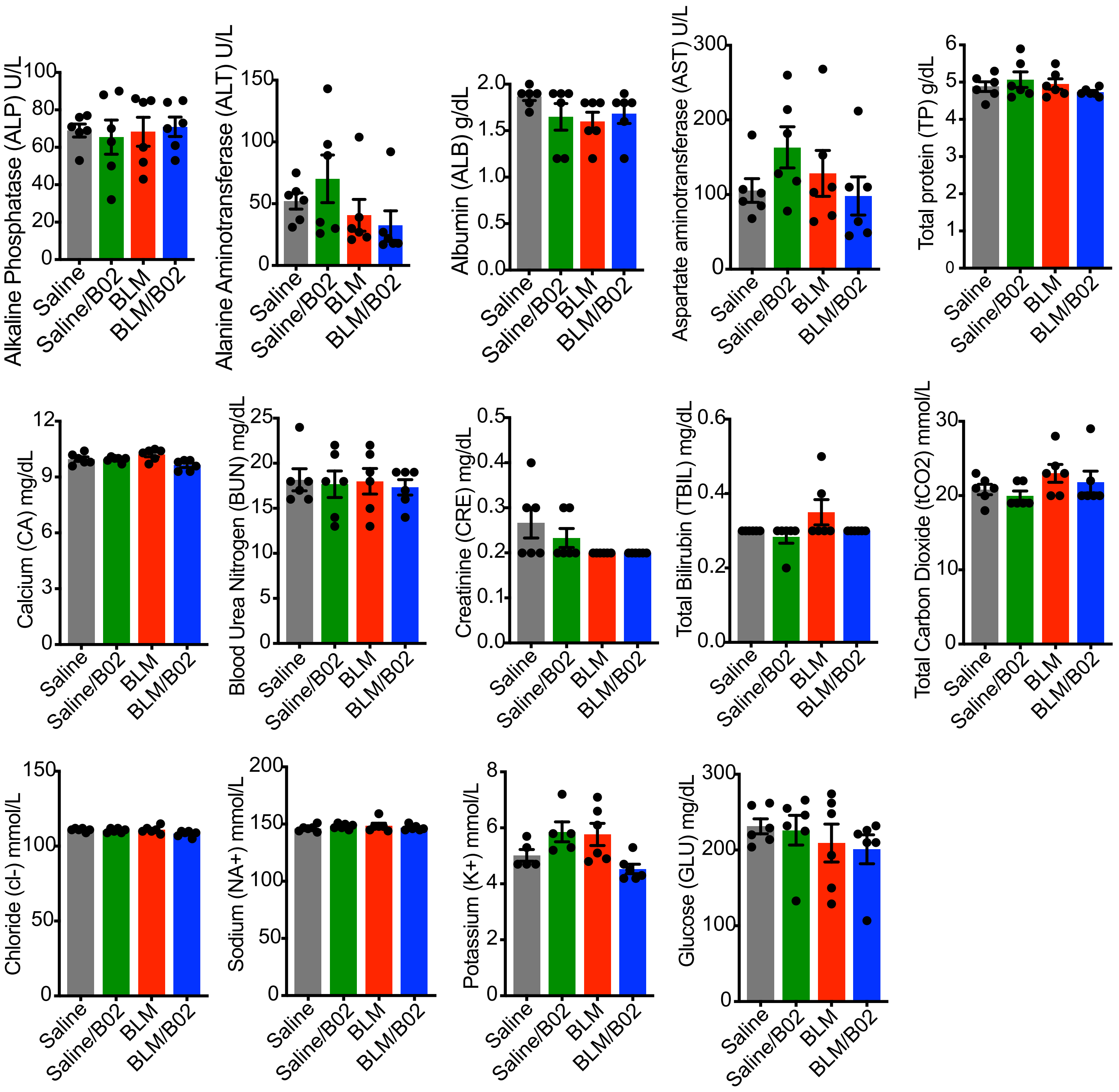

**Supplementary Figure S11. Rad51 inhibition does not exhibit any demonstrable impact on the liver or kidney of mice.** Male and female C57BL/6 mice were treated according to the procedures described in the Methods and in Figure 7a. Blood samples were collected on day 23 from the facial vein of unanesthetized animals using lithium heparin tubes. The serum concentrations of the specified parameters were analyzed with a Piccolo Xpress Chemistry Analyzer. Data are presented as mean ± SEM of n = 6.

**Supplementary Figure S12**

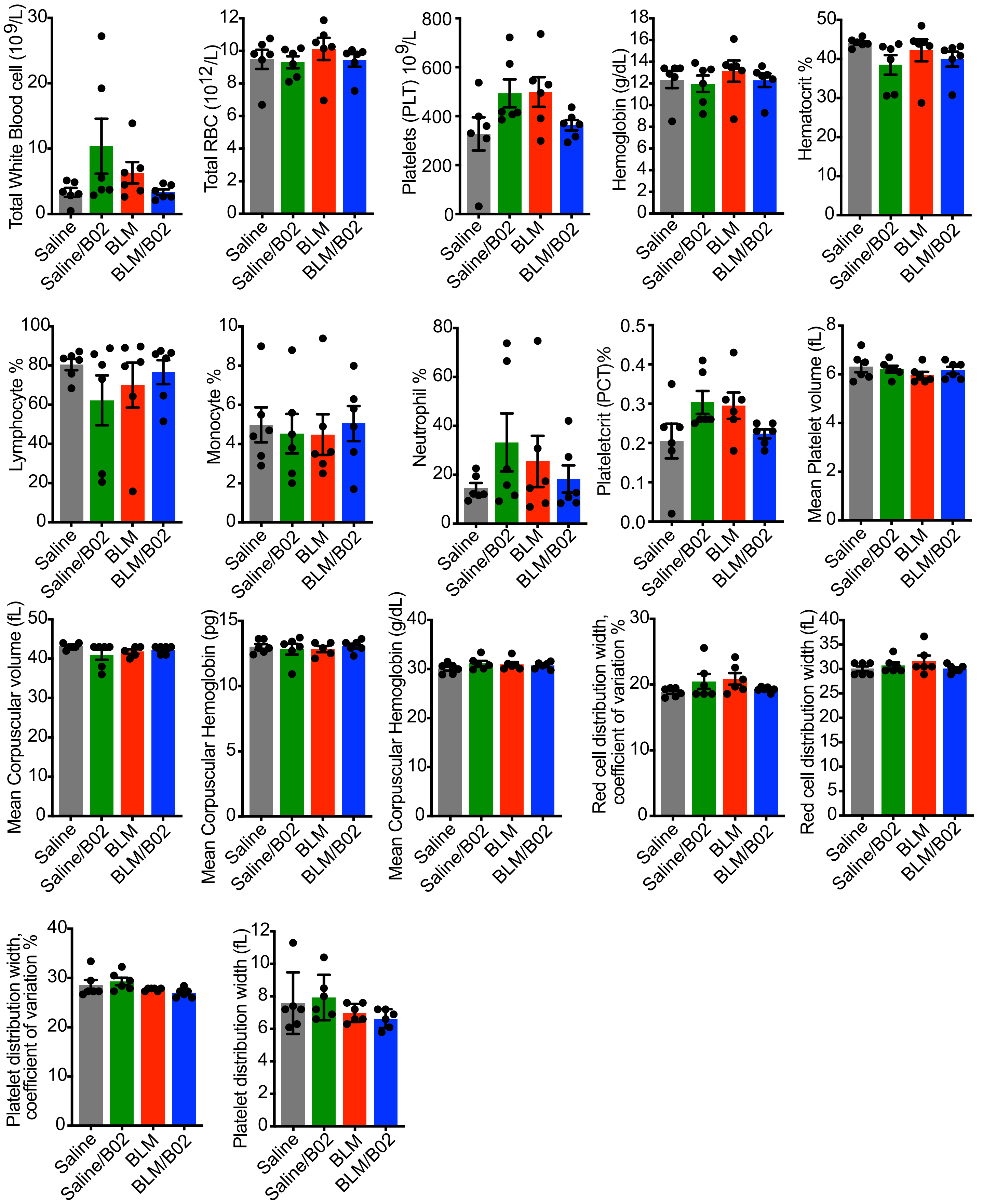

**Supplementary Figure S12. B02 treatment does not impair the recruitment of inflammatory cells.** C57BL/6 mice were treated as described in Methods and Figure 7a. On day 23, blood samples were collected in EDTA. Quantification of inflammatory cells and additional blood parameters was performed utilizing a VetScan HM5 Analyzer. Data are presented as mean ± SEM for n = 6.

**Table S1**

| REAGENT or RESOURCE | SOURCE | IDENTIFIER |
|---|---|---|
| **Antibodies** | | |
| Rabbit polyclonal anti-RAD51 Antibody | ThermoFisher | Cat# PA5-27195 |
| Cleaved Caspase-3 | Cell Signaling Technology | Cat# 9661S |
| Cytochrome c | Cell Signaling Technology | Cat# 4272S |
| VDAC | Cell Signaling Technology | Cat# 4866S |
| Donkey anti-goat IgG | Santa Cruz Biotechnology | Cat# sc-2020 |
| Goat anti-Rabbit IgG | Jackson ImmunoResearch | Cat# 111-035-003 |
| Goat anti-mouse IgG | Jackson ImmunoResearch | Cat# 115-035-003 |
| PUMA | ABclonal | Cat# A17138 |
| BAD | Cell Signaling Technology | Cat# 9292S |
| Glutaminase (GLS) | Novus Biologicals | Cat# NBP2-29940 |
| Histone H2A.X | Cell Signaling Technology | Cat# 2595S |
| Phospho-Histone H2A.X | Cell Signaling Technology | Cat# 9718S |
| Mouse monoclonal Anti- GAPDH antibody | Millipore | Cat# MAB374 |
| Goat polyclonal anti-Type I Collagen antibody | Southern Biotech | Cat# 1310-01 |
| Goat anti-mouse IgG (Alexa Fluor 488) | abcam | Cat# ab150113 |
| Goat anti-Rabbit IgG (Alexa Fluor 488) | abcam | Cat# ab150081 |
| Goat anti-Rabbit IgG (Alexa Fluor 594) | abcam | Cat# ab150080 |
| Donkey anti-Goat IgG (Alexa Fluor 488) | ThermoFisher | Cat# A-11055 |
| Anti-BAK antibody | abcam | Cat# ab32371 |
| Anti-BAX antibody | abcam | Cat# ab53154 |
| Anti-p53 (acetyl K120) antibody | abcam | Cat#AB78316 |
| Anti-alpha smooth muscle Actin antibody | abcam | Cat# ab7817 |
| Anti-alpha smooth muscle Actin antibody | abcam | Cat# ab124964 |
| CTHRC1 | Santa Cruz Biotechnology | Cat# sc-293270 |
| Rabbit monoclonal anti-CTGF antibody | Cell Signaling Technology | Cat# 86641S |
| Rabbit polyclonal anti-phospho-SMAD3 antibody | [1] | N/A |
| Rabbit monoclonal anti-SMAD3 antibody | Abcam | Cat# ab40854 |
| Mouse monoclonal anti-αSMA antibody | Sigma-Aldrich | Cat# A5228 |
| Rabbit polyclonal anti-Fibronectin antibody | Sigma-Aldrich | Cat# F3648 |
| Rabbit polyclonal anti-SMAD2 antibody | Abcam | Cat# ab63576 |
| Rabbit polyclonal anti-phospho-Smad2 antibody | [1] | N/A |
| Rabbit polyclonal anti-Akt antibody | Cell Signaling Technology | Cat# 9272 |
| Rabbit polyclonal anti-phospho-Akt (Ser473) antibody | Cell Signaling Technology | Cat# 9271 |
| Rabbit polyclonal anti-p70 S6 Kinase antibody | Cell Signaling Technology | Cat# 9202 |
| Rabbit polyclonal anti-phospho-p70 S6 Kinase (Thr389) antibody | Cell Signaling Technology | Cat# 9205 |
| Rabbit monoclonal anti-Phospho-4E-BP1 antibody | Cell Signaling Technology | Cat# 2855S |
| Rabbit monoclonal anti-4E-BP1 antibody | Cell Signaling Technology | Cat# 9644S |
| **Biological Samples** | | |
| Normal and IPF lung fibroblast | This manuscript and [2] | U of Pittsburgh IRB# 970946, WashU IRB #201103213 |
| **Chemicals, Peptides, and Recombinant Proteins** | | |
| DMEM (1X) + GlutaMAX | Gibco | Cat# 10569044 |
| B-27 Supplement (50X), serum free | Gibco | Cat# 17504044 |

| | | |
|---|---|---|
| Normal Donkey Serum | SouthernBiotech | Cat# 0030-01 |
| CTS Neurobasal Medium | Gibco | Cat# A1371201 |
| Hibernate-A Medium | Gibco | Cat# A1247501 |
| Seahorse XF DMEM assay medium | Agilent technologies | Cat# 103680-100 |
| Seahorse XF base medium | Agilent technologies | Cat# 103335-100 |
| UltraPure Low Melting Point Agarose | Invitrogen | Cat# 16520100 |
| Krebs-Ringer Solution, bicarbonate-buffered | Thermo Scientific | Cat# J67591.AP |
| Fetal Bovine Serum | HyClone Laboratories | Cat# SH3007103 |
| Penicillin-Streptomycin | Life Technologies | Cat# 15070063 |
| Glutamine | Agilent | Cat# 103579-100 |
| Protease Inhibitor Cocktail | Roche | Cat# 11836153001 |
| Recombinant Human TGFβ | R&D Systems | Cat# 240-B |
| Recombinant mouse TNFα | Biolegend | Cat# 575202 |
| Recombinant human TNFα | Biolegend | Cat# 570102 |
| LY294002 | Sigma-Aldrich | Cat# L9908 |
| MK2206 | Selleckchem | Cat# S1078 |
| Rapamycin | Selleckchem | Cat# S1039 |
| SB431542 | Tocris | Cat# 161410 |
| Lipofectamine 3000 | Thermo Fisher | Cat# L3000001 |
| U0126 | Promega | Cat# V1121 |
| B02 | Selleckchem | Cat# S8434 |
| Dimethyl sulfoxide | Sigma-Aldrich | Cat# D8418 |
| Phosphate Buffered Saline | Fisher bioreagents | Cat# BP399-1 |
| Bleomycin | Hikma | Cat# NDC 00143-9240-01 |
| **Critical Commercial Assays** | | |
| Glutamine Colorimetric Assay Kit | BioVision | Cat# K556 |
| RNeasy Plus Mini Kit | Qiagen | Cat# 74136 |
| PicoProbe Lactate Assay Kit | BioVision | Cat# K638-100 |
| Maxima Reverse Transcriptase | Life Technologies | Cat#EP0741 |
| Hydroxyproline Assay Kit | Sigma-Aldrich | Cat#MAK008 |
| Agilent Seahorse XF Cell Mito Stress Test Kit | Agilent Technologies | Cat#103015-100 |
| Agilent Seahorse XF Glycolysis Stress Test Kit | Agilent Technologies | Cat#103020-100 |
| ATP Colorimetric/Fluorometric Assay Kit | BioVision | Cat# K354 |
| NADP/NADPH Quantitation Kit | BioVision | Cat# K347-100 |
| Mitochondrial Transition Pore Assay Kit | Invitrogen | Cat# I35103 |
| WST-8 Assay Kit (Cell Proliferation) | abcam | Cat#ab65475 |
| **Oligonucleotides** | | |
| siRNA targeting human RAD51 | Santa Cruz Biotechnology | Cat# sc-36361 |
| siRNA targeting human SMAD2 | Santa Cruz Biotechnology | Cat# sc-38374 |
| siRNA targeting human SMAD3 | Santa Cruz Biotechnology | Cat# sc-38376 |
| Control siRNA | Santa Cruz Biotechnology | Cat#sc-37007 |
| Primers for all quantitative reverse transcription polymerase chain reaction (RT-qPCR), please refer to the Table S2 | This paper | N/A |
| **Software and Algorithms** | | |
| ImageJ | [3] | https://imagej.nih.gov/ij/ |
| GraphPad Prism 10.6 | | https://www.graphpad.com/scientific-software/prism/ |
| BioRender | | https://biorender.com |

**Table S2.**

**Primer for human qRT-PCR**

| Gene | Forward | Reverse |
|---|---|---|
| RAD51 | CAACCCATTTCACGGTTAGAGC | TTCTTTGGCGCATAGGCAACA |
| COL1A1 | GAGGGCCAAGACGAAGACATC | CAGATCACGTCATCGCACAAC |
| CTGF | GTCCAGCACGAGGCTCA | TCGCCTTCGTGGTCCTC |
| FN | TGTCAGTCAAAGCAAGCCCG | TTAGGACGCTCATAAGTGTCACCC |
| ACTA2 | GACAATGGCTCTGGGCTCTGTAA | CTGTGCTTCGTCACCCACGTA |
| TBP | GCCCGAAACGCCGAATATAATC | GTCTGGACTGTTCTTCACTCTTGG |

**Primer for mouse qRT-PCR**

| Gene | Forward | Reverse |
|---|---|---|
| Col1a1 | ATCTCCTGGTGCTGATGGAC | ACCTTGTTTGCCAGGTTCAC |
| Ctgf | CACAGAGTGGAGCGCCTGTTC | GATGCACTTTTTGCCCTTCTTAATG |
| Fn | TGACAACTGCCGTAGACCTG | ATCTAGCGGCATGAAGCACT |
| Tbp | GAAGTTCCCTATAAGGCTGGAAG | AGGAGAACAATTCTGGGTTTGA |
